# UBR5 is a Novel E3 Ubiquitin Ligase involved in Skeletal Muscle Hypertrophy and Recovery from Atrophy

**DOI:** 10.1101/592980

**Authors:** RA Seaborne, DC Hughes, DC Turner, DJ Owens, LM Baehr, P Gorski, EA Semenova, OV Borisov, AK Larin, DV Popov, EV Generozov, H Sutherland, II Ahmetov, JC Jarvis, SC Bodine, AP Sharples

**Author notes:** These authors contributed equally to the work (RA Seaborne, DC Hughes, DC Turner, DJ Owens, LM Baehr).

## Abstract

We aimed to investigate a novel and uncharacterised E3 ubiquitin ligase in skeletal muscle atrophy, recovery from atrophy/injury, anabolism and hypertrophy. We demonstrated an alternate gene expression profile for UBR5 versus well characterised E3-ligases, MuRF1/MAFbx, where after atrophy evoked by continuous-low-frequency electrical-stimulation in rats, MuRF1/MAFbx were both elevated yet UBR5 was unchanged. Furthermore, after recovery of muscle mass post tetrodotoxin (TTX) induced-atrophy in rats, UBR5 was hypomethylated and increased at the gene expression level, while a suppression of MuRF1/MAFbx was observed. At the protein level, we also demonstrated a significant increase in UBR5 after recovery of muscle mass from hindlimb unloading in both adult and aged rats, and after recovery from atrophy evoked by nerve crush injury in mice. During anabolism and hypertrophy, UBR5 gene expression increased following acute loading in three-dimensional bioengineered mouse muscle *in-vitro*, and after chronic electrical-stimulation-induced hypertrophy in rats *in-vivo*, without increases in MuRF1/MAFbx. Additionally, UBR5 protein abundance increased following functional overload-induced hypertrophy of the plantaris muscle in mice and during differentiation of primary human muscle cells. Finally, in humans, genetic association studies (>700,000 SNPs) demonstrated that the A alleles of rs10505025 and rs4734621 SNPs in the UBR5 gene were strongly associated with larger cross-sectional area of fast-twitch muscle fibres and favoured strength/power versus endurance/untrained phenotypes. Overall, we suggest that UBR5 is a novel E3 ubiquitin ligase that is inversely regulated to MuRF1/MAFbx, is epigenetically regulated, and is elevated at both the gene expression and protein level during recovery from skeletal muscle atrophy and hypertrophy.

**Key Points:**

- We have recently identified that a HECT domain E3 ubiquitin ligase, named UBR5, is altered epigenetically (via DNA methylation) after human skeletal muscle hypertrophy, where its gene expression is positively correlated with increasing lean leg mass after training and retraining.
- In the present study we extensively investigate this novel and uncharacterised E3 ubiquitin ligase (UBR5) in skeletal muscle atrophy, recovery from atrophy and injury, anabolism and hypertrophy.
- We demonstrated that UBR5 was epigenetically via altered DNA methylation during recovery from atrophy.
- We also determined that UBR5 was alternatively regulated versus well characterised E3 ligases, MuRF1/MAFbx, at the gene expression level during atrophy, recovery from atrophy and hypertrophy.
- UBR5 also increased at the protein level during recovery from atrophy and injury, hypertrophy and during human muscle cell differentiation.
- Finally, in humans, genetic variations of the UBR5 gene were strongly associated with larger fast-twitch muscle fibres and strength/power performance versus endurance/untrained phenotypes.

## Introduction

Proteolysis is the breakdown of proteins within cells. The ubiquitin-proteasome is the main system involved in protein turnover (Rock *et al.*, 1994; Cao *et al.*, 2005). In skeletal muscle, this system increases protein degradation as it reduces in size (reviewed in (Bodine & Baehr, 2014)) and remodels (Yang *et al.*, 2006; Gomes *et al.*, 2012; Slimani *et al.*, 2012). Ubiquitin-mediated proteolysis operates via distinctive enzymatic tagging to an area of a polypeptide chain using ubiquitin, securing the substrate proteins fate for degradation in the proteasome (Glickman & Ciechanover, 2002). Proteins tagged with ubiquitin are then recognised by the large 26S protease complex that degrades them into smaller peptides (Baumeister *et al.*, 1998). This ubiquitination tagging system is activated by enzymes triggered sequentially. The E1 enzyme activates ubiquitin, the E2 enzyme carries and conjugates ubiquitin to the E3 enzyme that then catalyses its binding to the designated protein substrate for degradation (reviewed in (Metzger *et al.*, 2012)). There are approximately 40 E2 and over 600 E3 ubiquitin ligases in the human genome making it an extremely complex and relatively uncharacterised system.

Over the past two decades, the E3 ligases in particular have received specific attention in skeletal muscle, namely due to the discovery of important E3’s that are selectively expressed in this tissue, including: Trim63, renamed muscle specific RING finger protein 1 (MuRF1); Fbxo32, renamed the F-box containing ubiquitin protein ligase atrogin-1 or Muscle Atrophy F-box (MAFbx/Atrogin-1) (Bodine *et al.*, 2001; Gomes *et al.*, 2001). As well as the more recently characterised; Fbxo30, renamed muscle ubiquitin ligase of SCF complex in atrophy-1 (MUSA1) (Sartori *et al.*, 2013), Parkin and ZNF216 (Hishiya *et al.*, 2006; Furuya *et al.*, 2014). MuRF1 contains a RING finger domain, whereas MAFbx/Atrogin-1 and MUSA1 have an F-box domain (part of the SCF family of E3s). RING fingers form the vast majority of all the E3 ligases. RING finger domains are zinc-binding motifs of 40-60 histidine and cysteine amino acid residues in a C_3_HC_4_ (RING) or C_3_H_2_C_3_ (RING-H2) formation (Bordone & Freemont, 1996). E3 RING finger proteins cause ubiquitin to be catalysed indirectly, acting as a scaffold that brings E2 proteins together with the substrate (Lorick *et al.*, 1999; Metzger *et al.*, 2012). Importantly, E3 ligases MuRF1 and MAFbx/Atrogin-1 have been demonstrated to increase rapidly at the gene expression level at around 48 hrs in the time course of atrophy in rodents following disuse (e.g. after denervation, hindlimb unloading, tetrodotoxin nerve exposure) increasing progressively up to 7-10 days (Bodine *et al.*, 2001; Gomes *et al.*, 2001; Clavel *et al.*, 2006; Moresi *et al.*, 2010; Fisher *et al.*, 2017). Furthermore, the expression of these genes increased in human muscle 48 hrs after spinal cord injury and 14 days after limb immobilisation (de Boer *et al.*, 2007; Urso *et al.*, 2007; Gustafsson *et al.*, 2010). MuRF1 and MAFbx/Atrogin-1 have also been shown to increase after starvation induced skeletal muscle atrophy (Gomes *et al.*, 2001; Lecker *et al.*, 2004) and are elevated in aging and cachexia (Gomes *et al.*, 2001; Lecker *et al.*, 2004; Clavel *et al.*, 2006; Costelli *et al.*, 2006; Raue *et al.*, 2007; Altun *et al.*, 2010). MuRF1 has been suggested to target myofibrillar proteins such as titin, nebulin, troponin1 (Witt *et al.*, 2005) and thick filament proteins such as myosin-binding protein C (MyBP-C) as well as myosin light chains 1 and 2 (MyLC1 and MyLC2) for degradation (Cohen *et al.*, 2009), whereas suggested MAFbx substrates include the myogenic regulatory factor, MyoD, Myogenin, eukaryotic translation initiation factor 3 subunit f (eIF3-f) (Tintignac *et al.*, 2005; Lagirand-Cantaloube *et al.*, 2008; Jogo *et al.*, 2009) and the intermediate filament proteins vimentin and desmin (Lokireddy *et al.*, 2012). Importantly, knock-down of MuRF1 and MAFbx/Atrogin-1 enzymes render mice partially resistant to denervation induced atrophy (Bodine *et al.*, 2001), and MuRF1 knock down also prevents atrophy following hindlimb unloading (HU) and glucocorticoid treatment (Bodine *et al.*, 2001; Labeit *et al.*, 2010; Baehr *et al.*, 2011). MuRF1 knock-down can help delay anabolic resistance and maintain muscle mass in aged rodents (Hwee *et al.*, 2014). Finally, MUSA1 knockdown using *in-vivo* electroporation of shRNA has been shown to spare muscle mass following denervation and prevent more severe atrophy observed in Smad4^−/−^ mice (Sartori *et al.*, 2013). Collectively, these data confirming an important function for these E3’s in skeletal muscle atrophy.

Together with RING finger domains, another key group of the E3 enzymes contain HECT (homologous to *E*6-AP carboxy-terminus) domains. The HECT domain of these E3 enzymes, unlike RING finger family members, directly binds activated ubiquitin to the target protein substrate. However, HECT domain E3 ligases remain relatively unstudied in skeletal muscle atrophy or remodelling. Interestingly, we have recently demonstrated that a HECT domain E3 ligase called UBR5 (aka. EDD1), previously uncharacterized in skeletal muscle, is an epigenetically regulated gene (identified using 850K DNA methylation arrays) hypomethylated after resistance exercise-induced muscle hypertrophy in humans (termed loading/training) (Seaborne *et al.*, 2018b). In this study, resistance exercise-induced hypertrophy was followed by complete cessation of exercise and a return of lean leg mass to pre-exercise baseline levels (unloading/detraining). In the same participants, a second period of resistance exercise-induced hypertrophy (reloading/retraining) ensued, in which the greatest increase in lean leg mass was observed. At the DNA level, we discovered that UBR5 became hypomethylated during periods of resistance exercise induced hypertrophy and returned to baseline levels upon exercise cessation. This oscillating methylation profile was inversely associated with changes in UBR5 expression, suggesting that the transcript was epigenetically regulated. Importantly, the largest increases in hypomethylation, gene expression and muscle mass were observed after later reloading/retraining. Therefore, the level of UBR5 gene expression was positively and strongly correlated with the increases in lean leg mass across after training and retraining (Seaborne *et al.*, 2018b). This data indicated that UBR5 was an E3 ligase associated with muscle hypertrophy rather than muscle atrophy. This was surprising given the well-defined role of the aforementioned E3 ligases in protein degradation, and therefore suggested that UBR5 may instead be important in muscle remodelling and recovery during changes in muscle mass. We therefore aimed to investigate the role of UBR5 more extensively during atrophy/injury and recovery from atrophy/injury as well as following anabolism, hypertrophy and muscle cell differentiation in order to start to characterise its role across these processes. We also aimed to further substantiate UBR5’s genetic role in muscle fibre size and physical performance using micro-array analysis in humans.

## Methods

### Tetrodotoxin (TTX)-induced atrophy and TTX cessation for recovery in-vivo

In new analysis of samples derived from previous work carried out by our group (Fisher *et al.*, 2017), we assessed UBR5 DNA methylation and gene expression following tetrodotoxin (TTX) induced atrophy and recovery of the tibialis anterior (TA) in rats. Briefly, under anaesthesia with isoflurane and buprenorphine, the common peroneal nerve in the left hindlimb of adult male Wistar rats was exposed to TTX using an implanted subcutaneous mini-osmotic pump (Mini Osmotic Pump 2002, Alzet, Cupertino CA, USA) in the scapular region, with a delivery tube leading to a silicone rubber cuff placed around the nerve. This delivered 0.5 μl/hr of TTX (350 μg/ml in sterile 0.9 % saline) so that the ankle dorsiflexor muscles (TA and extensor digitorum longus/EDL) were silenced but normal voluntary plantar-flexion was maintained for a period of 14 days (d). This evoked reductions in muscle weight of 7, 29 and 51% and fiber CSA of 18, 42, 69% at 3, 7 and 14 d respectively (Fisher *et al.*, 2017). The muscle was then allowed to recover after exhaustion in osmotic pump delivery of TTX to the peroneal nerve and allowing normal habitual physical activity to resume for a period 7 days post 14 d TTX atrophy (recovery), that resulted in a 52 and 63% increase after 7 d recovery of muscle weight and fiber CSA respectively versus the 14 d TTX-induced atrophy timepoint (Fisher *et al.*, 2017). RNA and DNA were isolated from the contralateral control (right) limb and from the TTX (left) limb at 3, 7, 14 d (n = 6 for 3/7/14 d, n = 4 recovery). Experimental procedures were conducted according to the permissions within a project license granted under the British Home Office Animals (Scientific Procedures) Act 1986. Wistar rats were sourced from registered breeding colonies at LJMU Life Sciences Support Unit and euthanasia was conducted by rising CO2 followed by cervical dislocation.

### Atrophy induced via continuous low frequency electrical stimulation in-vivo

In male Wistar rats (3 months of age) under anaesthesia with isoflurane and buprenorphine, a sterilised miniature implantable stimulator was inserted into the abdominal cavity and sutured to the abdominal wall, in a model previously described (Jarvis *et al.*, 1996). Briefly, in new experimentation, the electrodes were tunnelled subcutaneously and placed under (but not touching) the common peroneal nerve and sutured to the aponeurosis of the gastrocnemius muscle of the left hind limb to keep the electrodes in place, and allow activation of the TA and EDL via the common peroneal nerve. Animals were then allowed to recover from the implant procedure for 7 d before initiating a continuous low frequency 20 Hz stimulation for a further period of 7 d, turned on remotely using a stroboscope to transmit a sequence of light pulses through the skin. This resulted in an 12.64% reduction in muscle weight of the TA after 7 d in the stimulated versus the contralateral unstimulated right control limb (n = 6 muscle weight, Figure 1A), using methodology previously described (Jarvis *et al.*, 1996). Daily stimulation was conducted under no anaesthesia and amplitude of nerve stimulation was remotely adjusted under careful observation to provide motor-neurone activation without limb withdrawal or signs of distress. TA samples were taken from the stimulated and unstimulated contralateral limb at the end of the 7 d stimulation protocol. Experimental procedures were conducted according to permissions within a project license granted under the British Home Office Animals (Scientific Procedures) Act 1986. Wistar rats were sourced from registered breeding colonies at LJMU Life Sciences Support Unit and euthanasia was conducted by rising CO2 followed by cervical dislocation.

**Figure 1:**
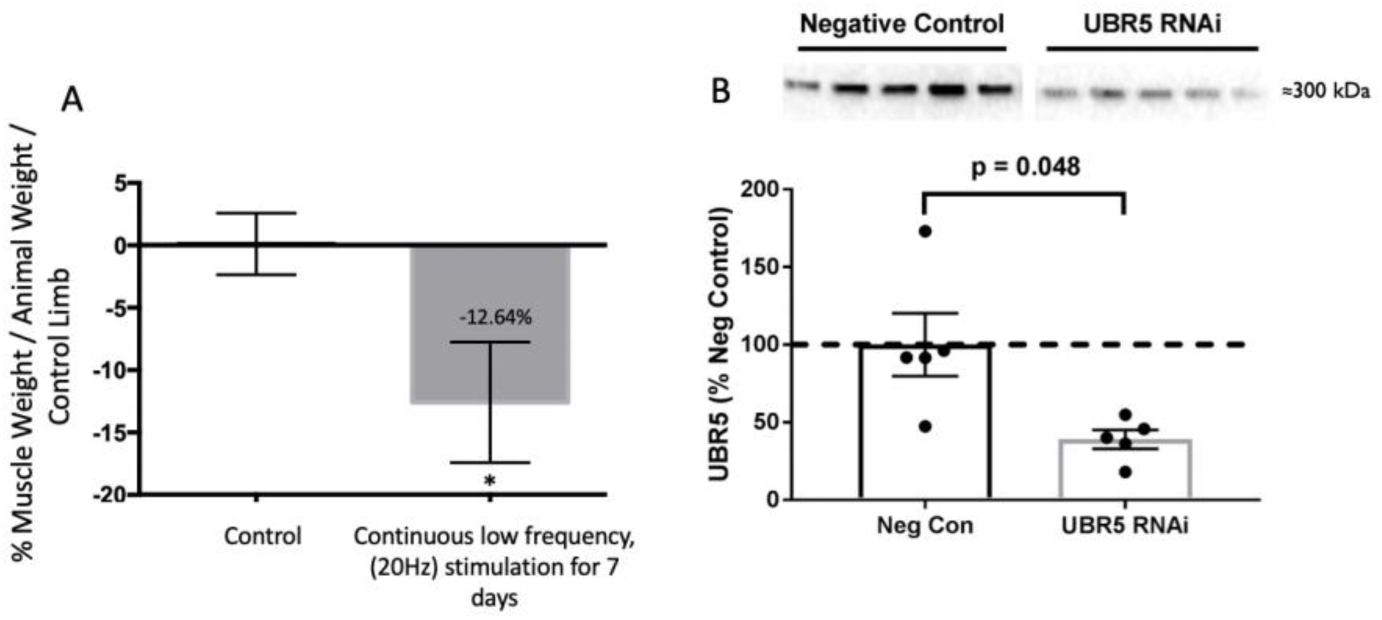
**A:** Reductions in muscle weight after 7 days of continuous low frequency (20Hz) electrical stimulation. TA weight was expressed relative to whole animal body mass to normalize for interindividual differences in animal size (n = 6). Then expressed as percentage change from the untreated contralateral control limb for each animal. **B**: UBR5 RNAi and negative control (neg con) plasmid injected into the TA muscle of C57/BL6 male mice (n = 5) after 7 days post electroporation. Western blots using UBR5 antibody (monoclonal, D6087; Cell Signalling Technology) demonstrate a significant reduction (p value depicted on figure) in UBR5 protein, validating the antibody for use in skeletal muscle.

### Hindlimb suspension induced-atrophy and recovery in adult and aged animals in-vivo

The response of hindlimb muscles to unloading and recovery was studied in 9-month (adult) and 29-month (aged) male Fischer 344-Brown Norway (F344BN) rats obtained from the National Institute of Aging. Using the same samples obtained from previous published experiments (Baehr *et al.*, 2016), we undertook new analysis of UBR5 protein abundance. Briefly, hindlimb unloading (HU) was achieved by tail-suspension as previously described (Thomason *et al.*, 1987; Baehr *et al.*, 2016), with the rats attached to a swivel (360° rotation) mounted plastic bar at the top of the cage in a ∼30° head-down tilt position for a period of 14 d. Animals were released from the tail suspension on day 15 at which time the rats were individually housed and allowed unrestricted cage activity for 3, 7, or 14 days (n = 6/ group). Muscle from the medial gastrocnemius (MG) was removed and lysed for protein extraction at baseline, 14 d after HU and at 3, 7 and 14 d of reloading/recovery (REL). During tissue collection, animals were anesthetized with isoflurane on completion of tissue removal, rats were euthanized by exsanguination. All animal procedures were approved by the Institutional Animal Care and Use Committee at the University of California, Davis, USA.

### Nerve crush injury, atrophy and recovery in-vivo

Targeted nerve crush in the lower limb muscles of the right leg was accomplished via acute 10 second crushing of the sciatic nerve in the midthigh region using forceps. The procedure was completed under isoflurane anaesthesia (3% inhalation) with the use of aseptic surgical techniques. Male mice (C57BL/6; 12 weeks old) purchased from Charles River Laboratories were given an analgesic (buprenorphine, 0.1 mg/kg) immediately and for 48 hours following the surgery and returned to their cage following recovery. Following completion of the appropriate time period (3, 7, 14, 21, 28, 45 and 60 days/d; n = 6/ group), mice were anesthetized with isoflurane, and the gastrocnemius complex (GSTC) muscles were excised, weighed, frozen in liquid nitrogen, and stored at −80°C for later analysis. On completion of tissue removal, mice were euthanized by exsanguination. Prior to tissue collection a sciatic nerve test was applied via a single electrical pulse to the sciatic nerve through electrodes to observe a twitch response in the hindlimb muscles. Between 21 and 60 days all animals displayed a twitch, suggesting that neural activity to the hindlimb muscles had returned at these time points. A separate untreated cohort of animals (n = 6) were used as relevant controls. All animal procedures were approved by the Institutional Animal Care and Use Committee at the University of Iowa, Iowa City, USA.

### Mechanical loading of bioengineered skeletal muscle in-vitro

Bioengineered mouse muscle was created using the C_2_C_12_ cell line as previously described by our group, using a fibrin self-assembling three-dimensional (3-D) model (Turner *et al.*, 2019a). Briefly, C2C12 cells were grown on T75’s to approx. 80% confluence whereby 90,000 cells/ml in 2 ml growth media (GM; 4.5 g/l glucose DMEM that included 4 mM L-glutamine, 10% hiFBS, 10% hiNBCS, supplemented with an additional 2 mM L-glutamine (Girven *et al.*, 2016),100 U/ml penicillin, 100 μg/ml streptomycin, 0.5 mg/ml 6-aminocaproic acid, 50 μM L-ascorbic acid, 50 μM L-proline) were seeded onto each sylgard coated (Sylgard 184 Elastomer Kit, Dow Corning, MI, USA) 35 mm culture dish containing a pre-polymerised (10 mins at room temperature and 1hr at 37 degrees) fibrin gel made up of 500 μl thrombin-solution (10 U/ml thrombin, 8 μl/ml aprotinin at 10 mg/ml) and 200 μl fibrinogen (20 mg/ml). GM was changed every 48 h until ∼90% confluency was attained. Media was then switched to differentiation media (DM; 4.5 g/l glucose DMEM that includes 4 mM L-glutamine, 2% HS, supplemented with an additional 2 mM L-glutamine, 100 U/ml penicillin, 100 μg/ ml streptomycin, 1 mg/ml 6-aminocaproic acid, 50 μM L-ascorbic acid, 50 μM L-proline) for 48 hrs and then then switched to 7% serum (same media constituents as DM with 7% serum, 3.5% hiFBS/ 3.5% hiNBCS) maintenance media (MM) to promote myotube formation for a further 10 days. Bioengineered muscle constructs were then transferred to bioreactor chambers (TC-3 Ebers, Spain, see Figure 4bi) for subsequent loading which involved 5 exercises × 4 sets × 10 reps (at 0.4 mm/s with 90 s rest between sets, 3.5 mins rest between exercises, totalling a stretch regime of 60 mins) at 10% stretch (1.2 mm) at 37°C / 5% CO_2_, recently described in full detail (Kasper *et al.*, 2017; Turner *et al.*, 2019a). To identify representative gel morphology (seen in Figure 4bi) bioengineered muscle constructs were fixed and dehydrated in ice-cold methanol:acetone:TBS (25:25:50) for 15 mins then a further 15 mins in methanol:acetone (50:50) only. Following fixation, gels were permeabilised (0.2% Triton X-100) in 1 × TBS for 90 mins and incubated overnight (4°C) in 250 μl Phalloidin-FITC (Sigma Aldrich, P5282) at a concentration of 50 μg/ml. After overnight incubation, Phalloidin-FITC was aspirated and gels were washed 3 × TBS before adding 250 μl of DAPI stain (ThermoFisher, D1306) at a concentration of 300 μM for 90 mins to counterstain myonuclei. Myotubes (Phalloidin-FITC, Excitation: 495 nm, Emission: 513 nm) and myonuclei (DAPI, Excitation: 358 nm, Emission: 461 nm) were visualised under an inverted fluorescence microscope (Nikon, Eclipse Ti-S) and imaged (Nikon, NIS Elements) for morphological analysis. RNA was isolated 30 minutes post-stretch to assess UBR5 transcript expression versus static non-loaded controls.

### Intermittent high frequency chronic electrical stimulation induced hypertrophy in-vivo

In adult male Wistar rats (6 months of age), under anaesthesia with isoflurane and buprenorphine, a sterilised miniature implantable stimulator was inserted into the abdominal cavity and sutured to the abdominal wall with electrodes placed near to the common peroneal and tibial nerves. The muscles were electrically stimulated to ensure that the dorsiflexors worked against the plantar-flexors, and therefore the dorsi-flexors were ‘loaded’. The stimulation consisted of an intermittent regime of high frequency (100 Hz) stimulation once a day for 4 weeks (5 sets × 10 reps, each rep 2s with 2s rest between reps, and 2.5 minutes rest between sets). Daily stimulation was conducted under no anaesthesia and amplitude of nerve stimulation was remotely adjusted under careful observation to provide motor-neurone activation without limb withdrawal or signs of distress. Following 4 weeks of the above regime there was an average 14% increase in muscle weight and an 19% increase in fiber CSA in the TA muscle at the mid-belly as previously described (Schmoll *et al.*, 2018). DNA and RNA was isolated from TA muscle from this original study (Schmoll *et al.*, 2018) from both the stimulated and contralateral (unstimulated) control limbs (n = 4) at the end of the 4 weeks electrical stimulation protocol. Experimental procedures were conducted according to the permissions within a project license granted under the British Home Office Animals (Scientific Procedures) Act 1986. Wistar rats were sourced from registered breeding colonies at LJMU Life Sciences Support Unit and euthanasia was conducted by rising CO2 followed by cervical dislocation.

### Synergist ablation/functional overload (FO) in-vivo

Hypertrophy of the plantaris muscle was induced in male C57BL/6 mice (12 weeks old) by surgical removal of synergist muscles (entire soleus and over half of the medial and lateral gastrocnemius without damaging the plantaris neural-vascular supply) according to a previously validated method termed synergist ablation or functional overload (FO) (McCarthy *et al.*, 2011; Bodine & Baar, 2012; Jackson *et al.*, 2012). The same protein lysates at baseline (control), 1, 3, 10 and 14 days (d) following FO previously derived in (Baehr *et al.*, 2014), that demonstrated significant increases in muscle weight of 43 and 65% after 7 and 14 d respectively, were analysed in new experimentation for UBR5 protein abundance in the present study (n = 3 / group). C57BL/6 mice were purchased from Jackson Laboratories. Mice were anaesthetized with inhaled isoflurane. Following tissues collection, the mice were euthanized by exsanguination. All animal procedures were approved by the Institutional Animal Care and Use Committee at the University of California, Davis, USA.

### Human muscle biopsies, cell isolation and differentiation in-vitro

Human muscle cells were isolated from the vastus lateralis biopsies of 3 healthy young adult males (28 ± 11 yr, 73.00 ± 0.91 kg, 177.2 ± 4.3 cm, means ± SD) who were recreationally active but had never undergone a period of supervised chronic aerobic or resistance exercise. Full biopsy procedure and primary cell isolation procedures are provided in (Seaborne *et al.*, 2018a; Seaborne *et al.*, 2018b; Turner *et al.*, 2019a; Turner *et al.*, 2019b). Ethical approval was granted by the NHS West Midlands Black Country, UK, Research Ethics Committee as previously reported (Seaborne *et al.*, 2018b). Briefly, conchotome biopsies were immediately transferred to a sterile class II BSC in pre-cooled transfer media (TM; Hams F-10, 2% hi-FBS, 100 U/ml penicillin, 100 μg/ml streptomycin and 2.5 μg/ml amphotericin-B) and any connective/adipose tissue removed using a sterile scalpel on a petri dish in the same transfer media. Biopsies were then washed in sterile PBS (plus 100 U/ml penicillin, 100 μg/ml streptomycin and 2.5 μg/ml amphotericin-B), scissor minced in the presence of 0.05% Trypsin/0.02% EDTA and then titrated on a magnetic stirring platform at 37°C, with this procedure repeated twice. The supernatant following each of these procedures was then collected and pooled with horse serum at 10% of the total supernatant volume to neutralize the trypsin. The supernatants from each treatment were then pooled and centrifuged at 340 rcf for 5 minutes to generate a cell pellet. The supernatant was plated in a T25 (as a backup source of any cells remaining) and the cell pellet was resuspended on a separate T25, both in growth media/GM (GM; Ham’s F10 nutrient mix supplemented with 10% hi-NCS, 10% hi-FBS, 100 U/ml penicillin, 100 μg/ml streptomycin, 2.5 μg/ml amphotericin B and 5 mM L-glutamine). Once confluent cells were split onto larger T75’s to generate enough cells to perform differentiation experiments. Human derived muscle cells were then seeded onto gelatin coated 6 well plates, at a density of 9×10^5^ cells/ml in 2 ml of GM for 24 hrs prior to removal of GM, 3 × PBS washes and transfer to differentiation media (DM; Ham’s F10 nutrient mix supplemented with 2% hiFBS, 100 U/ml penicillin, 100 μg/ml streptomycin, 2.5 μg/ml amphotericin B and 5 mM L-glutamine) for a period of 7 days. Cells were lysed for protein at 0 hrs (30 minutes in DM), 72 hrs and 7 days after transfer to DM. All experiments were carried out below passage 10 to prevent senescence.

### Human micro-array methods and association analyses

The human association studies involved 357 Russian international-level athletes (age 24.0 ± 3.6 years; 192 males and 165 females), 84 physically active men and 173 controls (83 males and 90 females). The athletes were stratified into 4 groups according to type, intensity, and duration of exercise. The first group (n = 142) included long-distance endurance athletes (3-10 km runners, biathletes, 5-10 km skaters, cross-country skiers, marathon runners, 0.8-25 km swimmers, race walkers, and triathletes. The second group (n = 115) comprised of sprinters (50-100 m swimmers, 200 m kayakers and canoers, sprint cyclers, 100-400 m runners, 500-1000 m skaters and short-trackers). The third group (n = 73) included strength athletes (powerlifters and weightlifters). The fourth group comprised explosive power athletes with predominantly anaerobic energy production (arm-wrestlers, track and field jumpers, heptathletes / decathletes and throwers). There were 198 athletes classified as ‘elite’ (prize winners of major international competitions). There were 159 athletes classified as ‘sub-elite’ (participants in international competitions). Physically active men (n=84; age 33.1 ± 7.3 years, height 180.1 ± 6.2 cm, weight 81.0 ± 10.4 kg were involved in the muscle biopsy study. Russian controls were 173 unrelated citizens of Russia without any competitive sport experience. The athletes and controls were all Caucasians of Eastern European descent. The overall study was approved by the Ethics Committee of the Physiological Section of the Russian National Committee for Biological Ethics and Ethics Committee of the Federal Research and Clinical Centre of Physical-Chemical Medicine of the Federal Medical and Biological Agency of Russia. Written informed consent was obtained from each participant. The study complied with the guidelines set out in the Declaration of Helsinki and ethical standards in sport and exercise science research.

Molecular genetic analysis in all Russian athletes, physically active men and controls was performed with DNA samples obtained from leukocytes (venous blood). Four ml of venous blood were collected in tubes containing EDTA (Vacuette EDTA tubes, Greiner Bio-One, Austria). Blood samples were transported to the laboratory at 4°C and DNA was extracted on the same day. DNA extraction and purification were performed using a commercial kit according to the manufacturer’s instructions (Technoclon, Russia) and included chemical lysis, selective DNA binding on silica spin columns and ethanol washing. Extracted DNA quality was assessed by agarose gel electrophoresis. HumanOmni1-Quad BeadChips (Illumina Inc, USA) were used for genotyping of 1,140,419 SNPs in athletes and controls and HumanOmniExpress BeadChips (Illumina Inc, USA) were used for genotyping of > 700,000 SNPs in participants of the Muscle Fiber Study (84 physically active men). The assay required 200 ng of DNA sample as input with a concentration of at least 50 ng/μl. Exact concentrations of DNA in each sample were measured using a Qubit Fluorometer (Invitrogen, USA). All further procedures were performed according to the instructions of the Infinium HD Assay.

### Evaluation of muscle fiber composition and cross-sectional area

Samples of the vastus lateralis muscle of Russian athletes were obtained with the Bergström needle biopsy procedure under local anaesthesia with 1% lidocaine solution. After this procedure, serial cross-sections (sections cut after on another) (7 μm) were obtained from frozen samples using a microtome (Leica Microsystems, Wetzlar, Germany). The sections were thaw-mounted on Polysine glass slides, kept for 15 min at RT and incubated with in PBS (3 × 5 min). Then one of the serial sections was incubated at RT in primary antibody against a slow isoform of the myosin heavy chains and the second serial section against a fast isoform of the myosin heavy chains (M8421, 1:5000; M4276; 1:600, respectively; Sigma-Aldrich, USA) for 1 h and incubated in PBS (3 × 5 min). After this, the serial sections were incubated at RT in a secondary antibody conjugated with FITC (F0257; 1:100; Sigma-Aldrich) for 1 hr. The sections were then washed in PBS (3 × 5 min), placed in mounting media and covered with a cover slip. The image was captured using a fluorescent microscope Eclipse Ti-U (Nikon, Japan). All analyzed images contained > 100 fibers. An example of stained images for slow and fast fibres can be seen in Figure 5A). The ratio of the number of stained fibers to the total fiber number was calculated. Fibers stained in serial sections with antibodies against both slow and fast isoforms were considered as hybrid fibers. The cross-sectional area (CSA) of fast and slow fibers was evaluated using ImageJ software (NIH, USA).

### Strength measurement

Evaluation of strength in elite Russian weightlifters was assessed by their performance in snatch, and clean and jerk (best results in official competitions including Olympic Games, Europe and World Championships). The total weight lifted (in kg) was multiplied by the Wilks Coefficient (Coeff) to find the standard amount lifted normalised across all body weights. 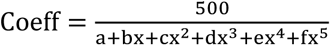, where x is the body weight of the weightlifter in kilograms. Values for males were: a = −216.0475144; b = 16.2606339; c =−0.002388645; d = −0.00113732; e = 7.01863E-06; f = −1.291E-08. Values for females were: a = 594.31747775582, b = −27.23842536447; c = 0.82112226871; d = −0.00930733913; e = 4.731582E-05; f = −9.054E-08.

### Gene expression by rt-qRT-PCR

Tissue from TTX atrophy/recovery, high frequency chronic intermittent electrical stimulation, continuous low frequency ‘overtraining’ stimulation and bioengineered muscle was homogenised in Tri-Reagent (Invitrogen, Loughborough, UK) for 45 seconds at 6,000 rpm x 3 (5 minutes on ice in between intervals) using a Roche Magnalyser instrument (Roche, Germany) and homogenization tubes containing ceramic beads (MagNA Lyser Green Beads, Roche, Germany). Human cells from differentiation experiments in monolayer were also lysed in Tri-Reagent for 5 minutes before being removed from the wells using a sterile cell scraper. RNA was then isolated from all these models using the manufacturer’s instructions for Tri-Reagent. In a one-step reaction (reverse transcription and PCR) qRT-PCR was performed using a QuantiFastTM SYBR^®^ Green RT-PCR one-step kit on a Rotorgene 3000Q, with a reaction setup as follows; 4.75 μl experimental sample (7.36 ng/μl totalling 35 ng per reaction), 0.075 μl of both forward and reverse primer of the gene of interest (100 μM stock suspension), 0.1 μl of QuantiFast RT Mix (Qiagen, Manchester, UK) and 5 μl of QuantiFast SYBR Green RT-PCR Master Mix (Qiagen, Manchester, UK). Reverse transcription initiated with a hold at 50°C for 10 minutes (cDNA synthesis) and a 5-minute hold at 95°C (transcriptase inactivation and initial denaturation), before 40-45 PCR cycles of; 95°C for 10 sec (denaturation) followed by 60°C for 30 secs (annealing and extension). Primer sequences for genes of interest and reference genes across mouse, rat and human samples are included in Table 1. All genes of interest and reference gene primers across species demonstrated no unintended targets via BLAST search and yielded a single peak after melt curve analysis conducted after the PCR step above. All relative gene expression was quantified using the comparative Ct (^ΔΔ^Ct) method (Schmittgen & Livak, 2008). For rat experiments that included gene expression (e.g. TTX atrophy-recovery, continuous low frequency electrical stimulation ‘overtraining’-induced atrophy and high frequency intermittent electrical stimulation induced hypertrophy), the contralateral control limb was used in ^ΔΔ^Ct equation as the calibrator condition. For bioengineered mouse muscle the pooled value from the static unloaded controls were used as the calibrator condition due to demonstrating low variation in Ct value (mean ± SD, 18.5 ± 0.8, 4.3% variation). Finally, for human cell differentiation the 0 hrs control sample within each subject was used as the calibrator. The average Ct value for the human cell differentiation experiments (RPL13a), rat (Polr2a) and mouse (Pol2rb) reference genes were also consistent across all samples and experimental conditions (human cell differentiation (mean Ct ± SD): 14.8 ± 0.5, rat TTX 21.8 ± 1.1, rat 20Hz continuous low frequency stimulation ‘overtraining’ 21.8 ± 1, rat intermittent high frequency 100Hz stimulation 23.2 ± 0.7, mouse bioengineered muscle 18.3 ± 0.7) with small variations of 3.5, 4.9, 4.7, 2.8, 3.9% respectively. The average PCR efficiencies of UBR5 (human cell differentiation 91.8%, rat TTX 91.7%, rat 20Hz continuous low frequency stimulation ‘overtraining’ 92.5%, rat intermittent high frequency 100Hz stimulation 90.4%, mouse bioengineered muscle 93.8%) were comparable with the human reference gene RPL13a (97.2%), rat reference Pol2ra gene (rat TTX 93.9%, rat 20Hz continuous low frequency stimulation ‘overtraining’ 93.5%, rat intermittent high frequency 100Hz stimulation 90.9%) and mouse Pol2rb gene (mouse bioengineered muscle 91.6%).

**Table 1.**
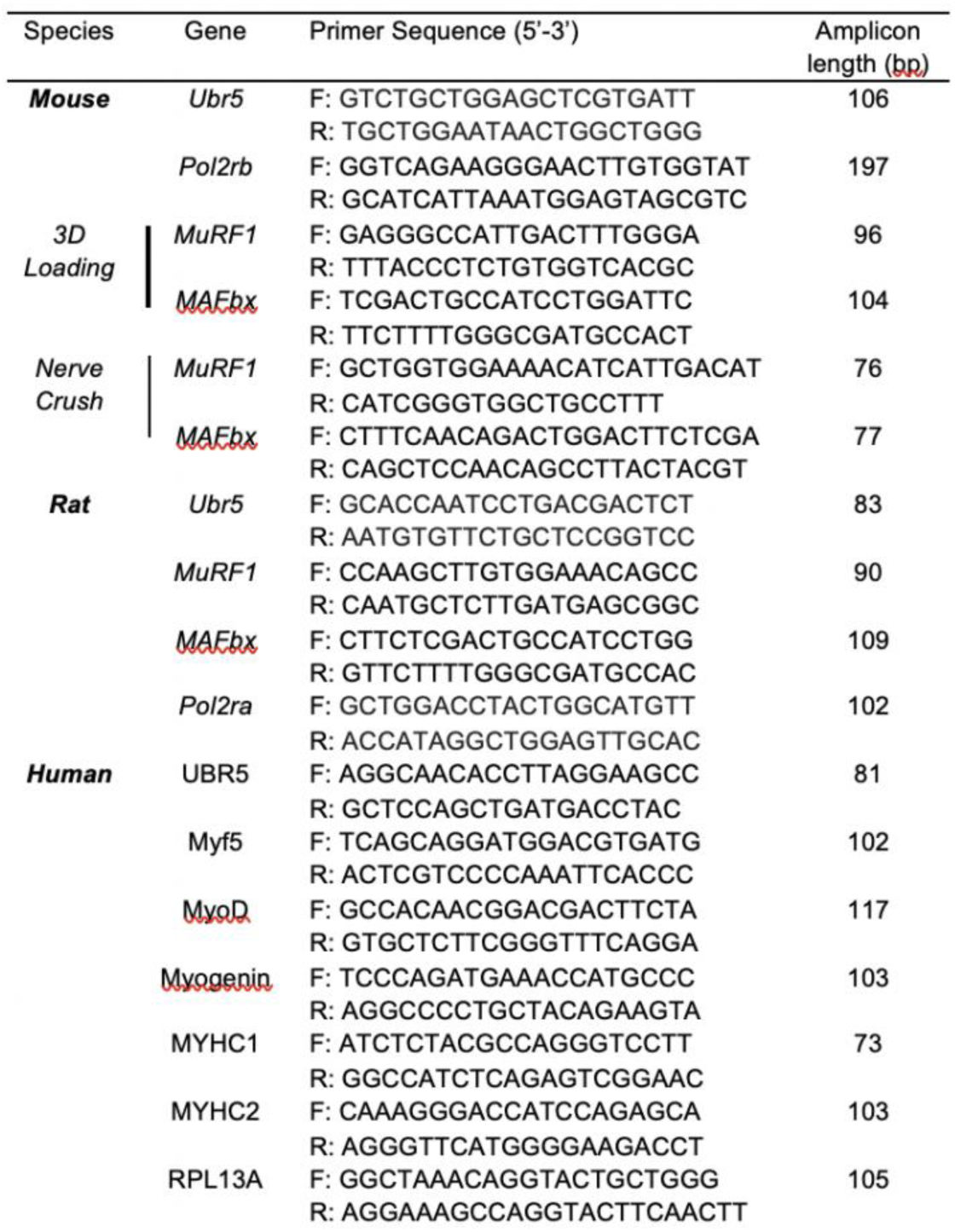
Gene expression primers.

Finally, for nerve crush injury model experiments, frozen muscle powder was homogenised using RNAzol RT reagent (Sigma-Aldrich, St Louis, USA) according to the manufacturer’s instructions. cDNA was synthesized using a reverse transcription kit (iScriptTm cDNA synthesis kit, Bio-rad, USA) from 1 μg of total RNA. 10 μl PCR reactions were setup as follows: 2 μl of cDNA, 0.5 μl (10 μM stock) forward and reverse primer, 5 μl of *Power* SYBR™Green master mix (Thermofisher Scientific, USA)) and 2 μl of RNA/DNA free water. Gene expression analysis was then performed by quantitative PCR (qPCR) on a Quantstudio 6 Flex Real-time PCR System (Applied Biosystems) using the UBR5 mouse primers in Table 1 (also used in the other mouse bioengineered muscle analysis). PCR cycling was as follows hold at 50°C for 5 minutes, 10-minute hold at 95°C, before 40 PCR cycles of; 95°C for 15 sec followed by 60°C for 1 minute (combined annealing and extension). Primer testing in these nerve crush samples yielded 89.04% efficiency for UBR5. As above, melt curve analysis at the end of the PCR cycling protocol yielded a single peak. For these experiments due to reference gene instability, after first assessing total RNA/mg tissue and then taking the ^Δ^Ct expression value, the mRNA was described relative to tissue weight, and then described as fold change relative to control muscles as previously described (Heinemeier *et al.*, 2009) an analysis used extensively by the group previously (Baehr *et al.*, 2016; Baehr *et al.*, 2017; Hughes *et al.*, 2018).

### DNA methylation by pyrosequencing

DNA samples from cells and tissue were isolated using Qiagen’s DNA easy blood and tissue kit (Manchester, UK) and bisulfite converted using Zymo Research EZMethylation Kit (Zymo Research, Irvine, CA, USA) as per manufacturer’s instructions. Prior to DNA isolation tissue samples were homogenized for 45 seconds at 6,000 rpm x 3 (5 minutes on ice in between intervals) in the lysis buffer (buffer ATL and protinase K) provided in the above DNA isolation kits using a Roche Magnalyser instrument and homogenization tubes containing ceramic beads (Roche, UK). UBR5 PCR primers and pyrosequencing primers were purchased from EpigenDX (Hopkinton, MA, USA), rat assay no: ADS5785 and mouse assay no: ADS5784, summarized in Table 2. PCR reactions were setup as follows; 3 μl of 10X PCR buffer (containing 15 mM MgCl^2^), plus an additional 1.8 μl of 25 mM MgCl^2^, 0.6 μl of 10 mM dNTPs, 0.15 μl HotStar Taq Polymerase (5 U/μl), 1 μl of bisulfite treated DNA and 0.6 μl of forward and reverse primer (10 μM). One primer was biotin-labelled and HPLC purified in order to facilitate purification of the final PCR product using streptavidin sepharose beads (see below). Following an initial denaturation incubation at 95°C for 15-min, for the Rat UBR5 assay; 45 cycles of denaturation at 95°C for 30 s; 51°C for 30 s (annealing), 68°C for 30 s (extension) were performed, with all PCR cycles followed by a final 5 minutes at 68°C. The mouse UBR5 assay had the same cycling parameters except for the annealing step that was conducted at 51°C for 30 s. PCR products were then bound to 2 μl Streptavidin Sepharose HP (GE Healthcare Life Sciences) in 40 μl Binding Buffer (Pyromark binding buffer, Qiagen, Manchester UK) and 25 μl Milli-Q-water. After which the immobilized PCR products were purified, washed, denatured with a 0.2 μM NaOH solution and rewashed using the Pyrosequencing Vacuum Prep Tool (Pyrosequencing, Qiagen, Manchester, UK), as per the manufacturer’s instructions. The denatured and washed PCR products were then placed in annealing buffer (Pyromark annealing & wash buffers, Qiagen, Manchester UK) with 0.5 μM sequencing primer at 75°C for two minutes on a heating block and the block then turned off for 10 minutes with the samples left in the block and then a further 10-15 minutes to cool at room temperature. Sequencing of the single stranded products yielded after PCR was conducted on a PSQ96 HS System (Qiagen, Manchester, UK) using Pyromark Gold Q24/Q96 reagents (Qiagen, Manchester, UK) following the manufacturer’s instructions. Methylation status of the individual CpG sites was identified as an artificial C/T SNP using QCpG software (Pyrosequencing, Qiagen, UK). Then % methylation of each CpG site was then calculated as a proportion of the methylated alleles divided by the sum of all methylated and unmethylated alleles, with the average methylation also then calculated across all CpG sites analysed within each gene. Non-CpG cytosines were included as internal controls for detection of incomplete bisulfite conversion. Furthermore, unmethylated and methylated DNA strands were included as controls after each PCR and bias testing was conducted by unmethylated control DNA mixed with *in vitro* methylated DNA at different ratios (0%, 5%, 10%, 25%, 50%, 75% and 100%). All supporting information for gene assays, including assay genomic sequence, bisulfite converted sequence and pyrosequencing dispensation order, CpG island locations, position from transcriptional start site and CpG ID numbers are given in Table 2.

**Table 2.**
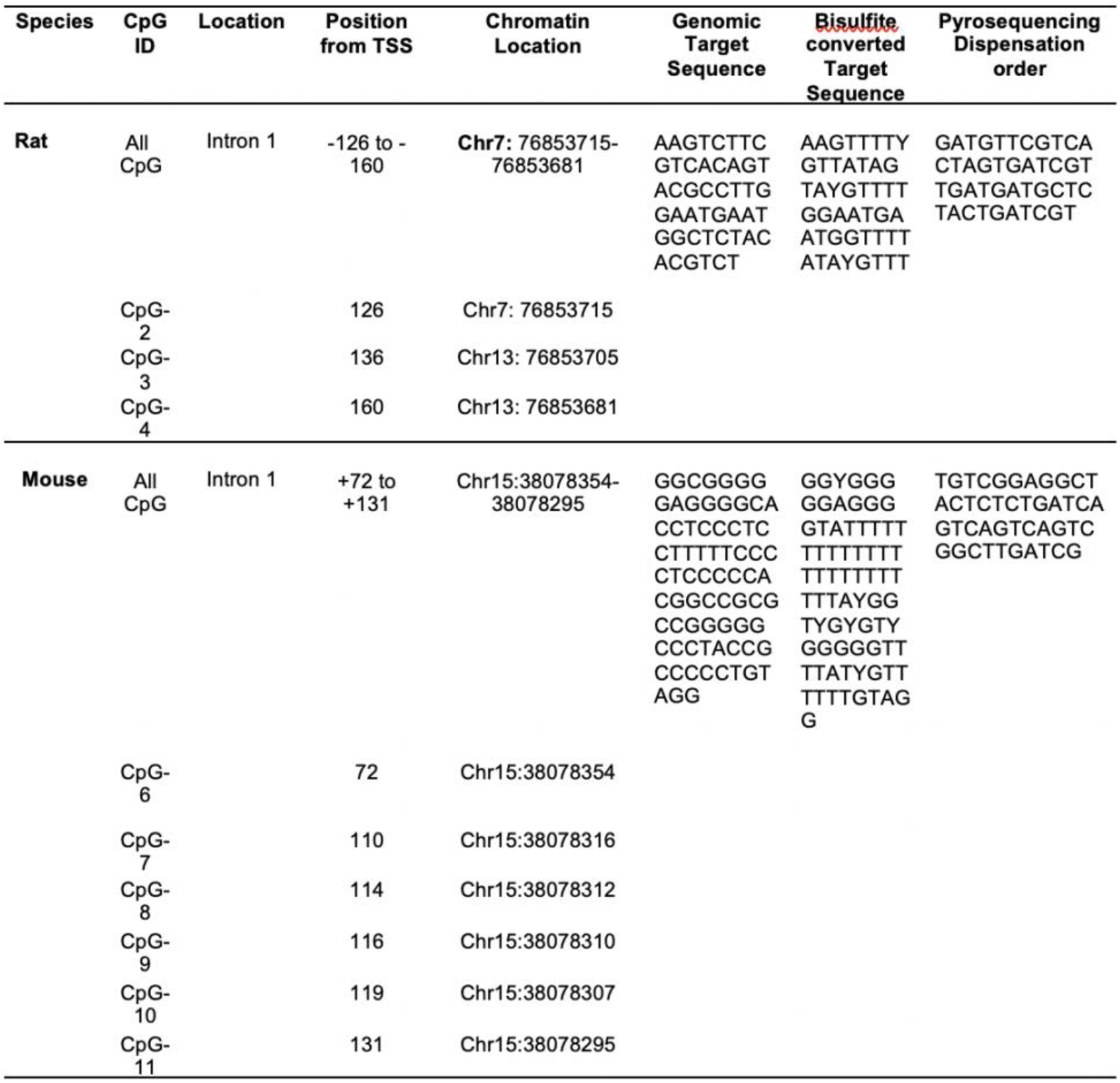
Description of targeted DNA methylation assays for loci specific pyrosequencing analysis of UBR5 in Rat and Mouse samples. The CpG loci location for the rat assay is based on Ensembl Gene ID (ENSRNOG00000006816), Ensembl transcript ID (ENSRNOT00000009115) and Rnor_6.0/rn6 genomic build. For the mouse assay it is based on Ensembl Gene ID (ENSMUSG00000037487), Ensembl transcript ID (ENSMUST00000226414) andGRCm38/mm10 genomic build.

### SDS page and western blotting

For UBR5 protein analysis from the hindlimb unloading, nerve crush and functional overload models, medial gastrocnemius (MG), gastrocnemius complex (GSTC) and plantaris muscle respectively were homogenized in sucrose lysis buffer (50 mM Tris pH 7.5, 250 mM sucrose, 1 mM EDTA, 1 mM EGTA, 1% Triton X 100, 50 mM NaF). The supernatant was collected following centrifugation at 8,000 *g* for 10 minutes and protein concentrations were determined using the Pierce™660 nm protein assay reagent (Thermo Scientific, Waltham, USA). Twelve micrograms of protein were subjected to SDS-PAGE on 4-20% Criterion TGX stain-free gels and transferred to polyvinylidene diflouride membranes. Membranes were blocked in 3% non-fat milk in Tris-buffered saline (TBS) with 0.1% Tween-20 (TBST) added for one hour and then probed with primary antibody overnight at 4°C (monoclonal UBR5 antibody, D6087; Cell Signalling Technology 1:500). Membranes were washed and incubated with HRP-conjugated secondary antibodies at 1:10,000 for one hour at room temperature. Immobilon Western Chemiluminescent HRP substrate was then applied to the membranes prior to image acquisition. Image acquisition and band quantification were performed using the Azure C400 System and Image Lab 6.0.1 software, respectively. Total protein loading of the membranes captured from images using stain-free gel technology (described above) was used as the normalization control for all blots for data from the hindlimb unloading, nerve crush and functional overload models. The same control lysates were run on two western blots to allow for quantification of UBR5 protein across multiple time points with the nerve crush injury analysis.

For human cell differentiation experiments, cell monolayers were lysed with trypsin and centrifuged at 18,000g to pellet the cells. The supernatant was discarded, and the remaining pellet was immediately frozen in liquid nitrogen. For the isolation of total protein, cell pellets were re-suspended in ice cold radioimmunoprecipitation assay (RIPA) buffer (Thermo Scientific) containing protease and phosphatase inhibitor cocktail (cOmplete Mini Protease Inhibitor Cocktail; Roche). The cell solution was kept on ice for ten minutes, following which samples were sonicated and vortexed before total protein content was determined by a bicinconinic acid assay (BCA) (Pierce, Thermo Fisher). Samples were then diluted to a final concentration of 1 μg/μl in 4x Laemmli buffer (Bio-Rad) and ultra-pure water and boiled for 10 minutes to denature proteins. For the determination of UBR5 abundance, equal amounts of protein (20 ug) were separated using SDS-PAGE on 6% gels at a constant voltage of 100 V per gel. Protein was then transferred onto a nitrocellulose membrane by wet transfer at 30 V overnight at 4^°C^ in Towbin buffer. Protein transfer was assessed by ponceau red staining (Sigma-Aldrich). Membranes were blocked with 5% bovine serum albumin (BSA) in TBST and then incubated overnight at 4^°C^ with a monoclonal UBR5 antibody (D6087; Cell Signalling Technology) and GAPDH (14C10; Cell Signalling Technologies) at a working concentration of 1:500 and 1:1000 in 5% BSA-TBST, respectively. Following 3 × 10 min TBST washes, membranes were washed in TBST 3 × 10 minutes prior to incubation in appropriate horse radish peroxidase-conjugated secondary antibody at room temperature for one hour. Chemiluminescent substrate was added to membranes and visualised in a Chemi-doc MP imaging system (Bio-Rad). Integrated density was determined for each band and all bands were normalized to GAPDH.

### Muscle specific knock-down of UBR5 for antibody validation

To confirm that the UBR5 antibody above (monoclonal UBR5 antibody, D6087; Cell Signalling Technology) used in the western blots was specific for UBR5. We use an RNAi system which was designed and purchase via Invitrogen (Massachusetts, *USA*). Specifically, the negative control RNAi plasmid was described previously (Ebert *et al.*, 2012) and encodes emerald green fluorescent protein (EmGFP) and a non-targeting pre-miRNA under bicistronic control of the CMV promoter in the pcDNA6.2GW/EmGFP-miR plasmid (Invitrogen). The UBR5 RNAi plasmid encodes EmGFP and an artificial pre-miRNA targeting mouse UBR5 under bicistronic control of the CMV promoter; it was generated by ligating the Mmi571982 oligonucleotide duplex (Invitrogen) into the pcDNA6.2GW/EmGFP-miR plasmid. The designed plasmids were incubated at 37°C overnight with TopOne Shot cells (Thermofisher Scientific cat no. C404010) on LB Agar Bacterial resistant plates (50 μg/ml Spectinomycin Sigma Aldrich cat no. S4014) in SOC medium (Invitrogen cat no. 15544-034), with isolated colonies then incubated/grown overnight in an orbital shaker at 37°C in falcon tubes. Bacteria where then pelleted at 6800 g for 1 min. Plasmid DNA f rom the bacteria was then prepared using a maxi prep kit (Nucleobond Xtra Maxi EF, Cat no. 740424.50 Machery-Nagel, USA) as per manufacturer’s instructions. The electroporation of DNA plasmids was performed as previously described (Ebert *et al.*, 2010; Hughes *et al.*, 2018). Briefly, after a 2 hr pre-treatment with 0.4 units/μl of hyaluronidase, 20 μg of mIR RNAi or negative control plasmid was injected into the TA muscle in C57/BL6 male mice (12-16 week-old, n = 5; purchased from Charles River Laboratories) and the hind limbs were placed between two-paddle electrodes and subjected to 10 pulses (20 msec) of 175 V/cm using an ECM-830 electroporator (BTX Harvard Apparatus). It has been previously demonstrated that the use of a pre-treatment with hydaluronidase allows for enhanced levels of transfected muscle without increased levels of muscle damage during the electroporation process (McMahon et al., 2001). The TA muscle was harvested for western blot analysis after 7 days post electroporation. On completion of tissue removal, mice were euthanized by exsanguination. All animal procedures were approved by the Institutional Animal Care and Use Committee at the University of Iowa, Iowa City, USA. We subsequently confirmed that UBR5 protein abundance was significantly reduced following injection of the UBR5 RNAi plasmid into basal skeletal muscle (Figure 1B).

### Statistics

TTX atrophy/recovery, FO induced hypertrophy and nerve crush models used a two-way ANOVA with an appropriate post-hoc correction (Fishers LSD). Human cell differentiation experiments were analysed using a one-way ANOVA with Fisher LSD post-hoc test. Those with an experimental group versus a relevant control were analysed using t-tests (chronic low frequency electrical stimulation induced atrophy, intermittent high frequency-electrical stimulation induced hypertrophy, loading of bioengineered muscle, UBR5 antibody validation). Biological repeat numbers (n) are specific in the methods sections or associated figure legends. Human microarray and association study data were analysed using PLINK v1.90, R (version 3.4.3), and GraphPad InStat (GraphPad Software, Inc., USA) software. Differences in phenotype between different genotype groups were analysed using Wald test (for three genotypes). Allele frequencies between groups of athletes and controls were compared by χ^2^ testing. *P* values ≤ 0.05 were considered statistically significant.

## Results

### UBR5 following skeletal muscle atrophy and recovery

Utilising a novel *in-vivo* model of disuse, whereby the common peroneal nerve of Wistar rats was exposed to tetrodotoxin (TTX) for 3, 7 and 14 days, we were able to successfully induce progressive skeletal muscle atrophy, as we previously described (Fisher *et al.*, 2017). Here, we reanalyzed RNA/DNA from these samples and identified that UBR5 gene expression significantly increased early in the time-course of atrophy (3 days, p = 0.04), coincident with significant hypomethylation of the UBR5 promoter (p = 0.004, Figure 2A). UBR5 gene expression remained elevated at 7 days following TTX induced atrophy (p = 0.04) but reduced later at 14 days, by which time the most extensive muscle mass loss occurred (51% loss in muscle weight), suggesting that increased UBR5 may be associated with early remodelling as a result of atrophy, but not later in the timecourse where the most extensive atrophy occurred. To further support and investigate these findings, we examined a secondary muscle mass loss stimulus *in-vivo*. We exposed Wistar rats to continuous low frequency electrical stimulation (20 Hz) for 7 days to mimic a model of over-use induced atrophy, which we successfully accomplished (12.64% reduction in muscle weight; Figure 1A). Interestingly, we identified that UBR5 did not significantly increase compared to control (p = N.S), whereas MuRF1 and MAFbx were significantly increased (Figure 2B; MuRF1 p = 0.01, MAFbx P = 0.03).

To examine the temporal pattern of UBR5 during recovery of skeletal muscle, we utilised our 14 d TTX atrophy model with an extended 7 day TTX cessation period to allow recovery from atrophy. This model successfully recovered skeletal muscle mass and CSA by 52 and 63% respectively (Fisher *et al.*, 2017). In this recovery model, we identified significant hypomethylation of UBR5 promoter (p = 0.04, Figure 2A) with an average increase in gene expression of UBR5 that did not quite reach significance (p = 0.1; Figure 2A & C). Importantly, MuRF1 and MAFbx were also significantly increased at the gene expression level at 3 and 7 days post TTX-induced atrophy (all p ≤ 0.01), as previously described in (Fisher *et al.*, 2017), yet these E3 ligases were upregulated to a greater extent (larger fold change) over the time course of atrophy than UBR5 (Figure 2C), and importantly were downregulated at the gene expression level during recovery. During recovery of muscle, UBR5 showed a trend for upregulation (1.91 fold) (albeit non-significantly p = 0.1) versus MuRF1 and MAFbx that were supressed below baseline levels (downregulated) that resulted in a reduction vs. the 14 d TTX atrophy timepoint (p = 0.08, Figure 2C). This pattern of UBR5 upregulation and MuRF1 and MAFbx suppression during muscle growth was also observed after chronic intermittent stimulation induced hypertrophy in rats (p = 0.02, described below in the results and depicted in Figure 4ci). Overall, these data suggest that UBR5 hypomethylation (significant) and an increase in gene expression (non-significant/p = 0.1) were associated with recovery from atrophy and muscle growth (Figure 2A & C).

It is worth noting that in the TTX model, only 7 days of recovery was sufficient to reverse a loss of approximately half the weight of the TA seen after 14d of TTX. Therefore, this could be an explanation for observing significant hypomethylation (that precedes gene expression) yet a non-significant average increase in UBR5 gene expression itself. To further investigate changes in UBR5 gene and protein expression, we used another well characterised model of disuse atrophy and recovery, hind-limb unloading (HU) and recovery/reloading. In this model we analysed the medial gastrocnemius (MG) of adult rats that has previously been characterised to demonstrate a full recovery of mass after 14 days following resumed habitual physical activity (reloading) post HU induced-atrophy (Baehr *et al.*, 2016). Furthermore, as aging results in reduced muscle mass and an impaired recovery from atrophy (Baehr *et al.*, 2016) versus adult rats, we also examined changes in UBR5 in response to HU atrophy and reloading (recovery) in aged rats. Indeed, following 14 days of hind-limb suspension in adult and aged rats, UBR5 protein increased early (3 days, albeit non-significantly) during atrophy but returned to baseline by 7-14 days in both adult and aged skeletal muscle (Figure 2D & E adult and aged respectively). Importantly, during recovery of the MG following hindlimb unloading, the largest increase in UBR5 protein was observed, demonstrating significant increases at 7 and 14 days post loading in young adult (p = 0.057 at 14 d) and aged rats (both p = 0.02 at 7 and 14 d, Figure 2D & E). Overall, these data suggest that UBR5 protein increased in abundance and was associated with early remodelling during atrophy, similarly to the mRNA responses observed in the TTX atrophy model, and importantly were associated with muscle growth and remodelling during the reloading/recovery period in both adult and aged rats.

**Figure 2.**
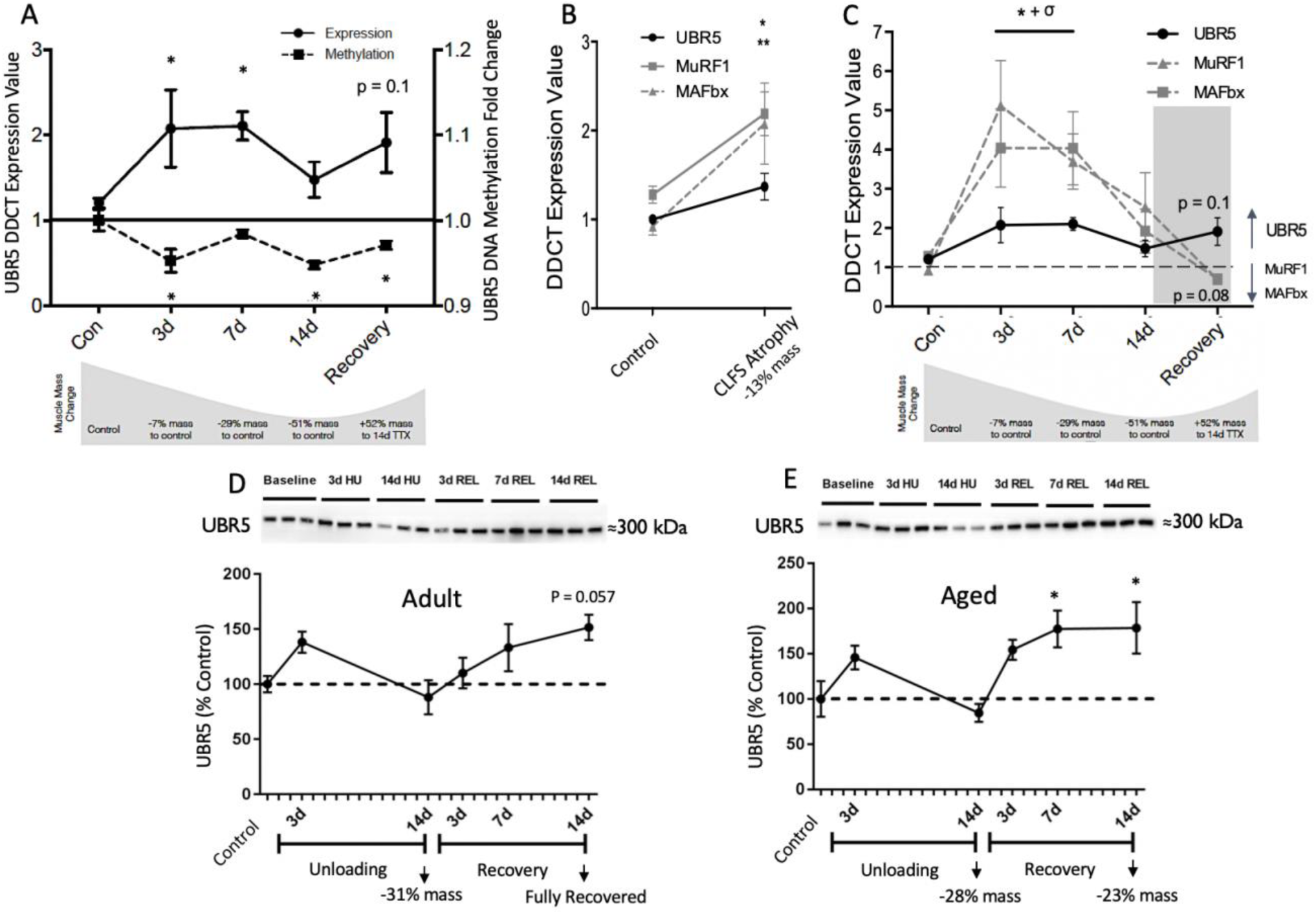
**A:** UBR5 DNA methylation and gene expression during Tetrodotoxin (TTX) atrophy and recovery in rat tibialis anterior (TA) (n = 6 for 3/7/14 days (d); n = 4 for recovery). Significant (*) increases in gene expression of UBR5 at 3 and 7 d. Significant (*) hypomethylation of the UBR5 promoter at 3 and 14 d of TTX and after 7 days recovery. **B:** No change in UBR5 gene expression compared with significant increases in MuRF1 (*) and MAFbx (* *) after 13% muscle mass atrophy as a result of ‘overtraining’ induced by continuous low frequency (20Hz) electrical stimulation (CLFS) (n = 3). **C:** UBR5 gene expression in comparison with MuRF1 and MAFbx taken from previously published data in the same conditions/samples (Fisher *et al.*, 2017) during TTX atrophy and recovery from atrophy (n = 6 for 3/ 7 / 14 d, n = 4 for recovery). All genes significantly increased early during atrophy at 3 (*) & 7 (σ) days, but with larger absolute fold changes in MuRF1/MAFbx vs. UBR5. During recovery MuRF1 and MAFbx were downregulated (MuRF1, p = 0.08 vs. 14 D) whereas UBR5 was upregulated (p = 0.1 vs. control). **D & E**: Protein abundance of UBR5 during hindlimb unloading (HU) induced atrophy and reloading/recovery (/REL) in the medial gastrocnemius muscle of adult (D) and aged (E) rats (n = 6/group). UBR5 was increased early after 3 d atrophy without significance. UBR5 then returned to baseline after 14 days of HU. Then in both adult (D) and aged (E) muscle UBR5 rapidly increased during recovery/REL resulting in significance (*) at 14 days in adult rats (p = 0.057), and both 7 and 14 days in aged rats (*). All graphs are means ± SEM. Significance at the level of p ≤ 0.05 unless otherwise stated.

### UBR5 after nerve crush injury

Because increased UBR5 was associated with recovery from atrophy we further sought to characterise its changes in a relevant muscle injury model of atrophy/recovery using a nerve crush approach. The sciatic nerve in the right lower limb of C57BL/6 mice was crushed resulting in denervation and subsequent reinnervation of the lower limb muscles by their original motoneurons. This model resulted in a significant reduction in weight of the gastrocnemius complex (GSTC) at 7, 14, 21 and 28 d (all p ≤ 0.05), the largest reduction (38% in muscle weight) occurring prior to reinnervation at 14 days post nerve crush injury (Figure 3A). The muscle fully recovered after 48 - 60 days (31 - 46 days after maximal atrophy) demonstrated by no significant changes vs. baseline controls (at 45 and 60 days, both p > 0.05). As with TTX and HU atrophy, UBR5 gene expression and protein abundance significantly increased early 3 days following nerve crush injury (Figure 3B gene expression, Figure 3D protein abundance, all P = 0.001). Furthermore, UBR5 gene expression dropped below baseline after 7 days (Figure 3B), whereas MuRF1 and MAFbx remained elevated (Figure 3C & D, albeit not significantly vs. baseline controls, they were significantly elevated vs. UBR5 at 7 days, both p = 0.01 for both MuRF1/MAFbx vs. UBR5). However, unlike recovery from TTX atrophy or HU during the recovery period, after 14 to 60 days of nerve crush injury, gene expression did not increase (Figure 3B) but significantly decreased in a similar fashion to MuRF1 and MAFbx (Figure 3C & D, all P = 0.001). However, importantly (and in an inverse relationship to UBR5 gene expression), protein abundance of UBR5 significantly increased over the time-course of recovery from nerve crush (Figure 3E) culminating in significant increases at 28 and 45 d (both P = 0.001), similar to the increase demonstrated above following recovery from HU.

**Figure 3.**
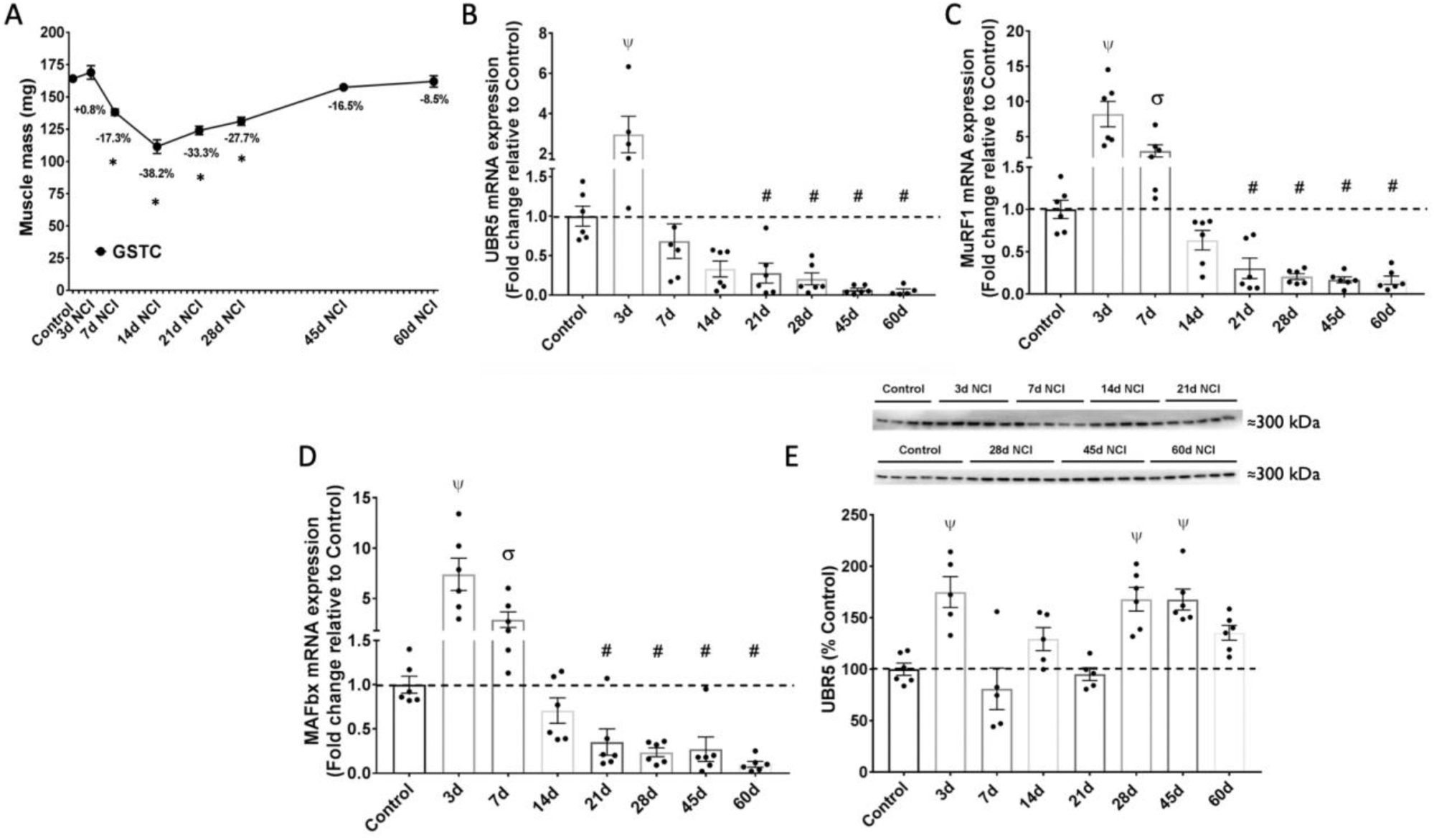
**A:** Muscle weight (mg) of the gastrocnemius complex muscles (GSTC) in mice after nerve crush injury (NCI) over a period of atrophy (3 −14 days/d) and recovery (14 d - 60 d) of muscle mass. The mice demonstrated a significant reduction in muscle weight of the GSTC at 7, 14, 21 and 28 d (all p ≤ 0.05 *) culminating in the largest average reduction of 38% in muscle weight at 14 days post nerve crush injury after which point the muscle fully recovered between 45 - 60 days (31 - 46 days after maximal atrophy) demonstrated by no significant changes vs. baseline controls at 45 and 60 days (both p > 0.05). **B, C, D:** UBR5 (B), MuRF1 (C) and MAFbx (D) significantly (Ψ p ≤ 0.001) increased at the gene expression level after 3 d post nerve crush injury and gene expression significantly (# p ≤ 0.001) decreased at 21, 28, 45-and 60-d post nerve crush injury. UBR5 returned towards baseline at 7 days whereas MuRF1 and MAFbx remained elevated at this timepoint post nerve crush injury (both p = 0.01 σ vs. UBR5 at 7 days, depicted on figure C & D). **E:** UBR5 protein abundance significantly increased early at 3 days post nerve crush injury, then returned to towards baseline after 7 and 14 d of atrophy and early during recovery at 21 d, before significantly increasing at 28 and 45 d during recovery (Ψ p ≤ 0.001). All graphs are means ± SEM. N = 5-6/group.

### UBR5 following skeletal muscle anabolism and hypertrophy

Our recently published data in humans suggests that UBR5 hypomethylation (via 850K DNA methylation arrays) and increased gene expression are associated with hypertrophy in humans following training and retraining (re-depicted in Figure 4A), with increased UBR5 gene expression positively correlating with increases in lean mass in humans (Seaborne *et al.*, 2018b). In the present study, we demonstrate that following an acute anabolic mechanical loading regime in mouse bioengineered muscle, there was also a significant increase (1.58 fold, p = 0.02) in UBR5 gene expression, as well as MAFBx (1.4 fold, P = 0.03) with a trend for increased expression of MuRF1 (1.77 fold, p = N.S) (all Figure 4Bi & Bii). Interestingly, this increase in UBR5 was similar to the fold change increase (1.71 fold) we reported previously, in humans after acute resistance exercise (Seaborne *et al.*, 2018b) (Figure 4Bi). In this previous paper (Seaborne *et al.*, 2018b), we identified that CpG methylation change of UBR5 did not change following the acute exercise stimulus (only after chronic training). We therefore examined promotor methylation of UBR5 in our mechanically loaded bioengineered mouse model to see if these findings translated across different models and species. In accordance with this, we identified that only 1 of the 6 CpG sites investigated in the UBR5 promoter were hypomethylated and the rest were hypermethylated following the acute anabolic loading stimulus, all non-significantly (data available on request). Further, in our original article, we showed an increase in gene expression following repeated/chronic resistance exercise (Seaborne *et al.*, 2018b), redrawn in Figure 4A). In an attempt to further support these findings, we reanalysed samples from an *in-vivo* rat model of chronic intermittent electrical stimulation induced hypertrophy, alongside a 14% increase in muscle weight of the TA (Schmoll *et al.*, 2018). Here we confirmed a similar trend as after human hypertrophy demonstrated previously ((Seaborne *et al.*, 2018b), redrawn in Figure 4A), with a significant increase in gene expression (Figure p = 0.02), and an average global reduction in methylation of the UBR5 promoter that did not reach significance (Figure 4Ci). Similarly, to our analysis following atrophy and recovery, we examined whether gene expression changes in UBR5 were similar or different to the other well-known E3 ubiquitin ligases, MuRF1 and MAFbx. Interestingly, while we observed a significant increase in UBR5 gene expression in the intermittent electrically stimulated hypertrophy model (Figure 4Ci), MuRF1 and MAFbx were unchanged in the same model of hypertrophy (Figure 4Cii). Therefore, all the E3 ligases were increased after acute mechanical loading but only UBR5 was increased, without an increase in MuRF1/MAFbx, at the gene expression level after chronic stimulation induced hypertrophy.

**Figure 4.**
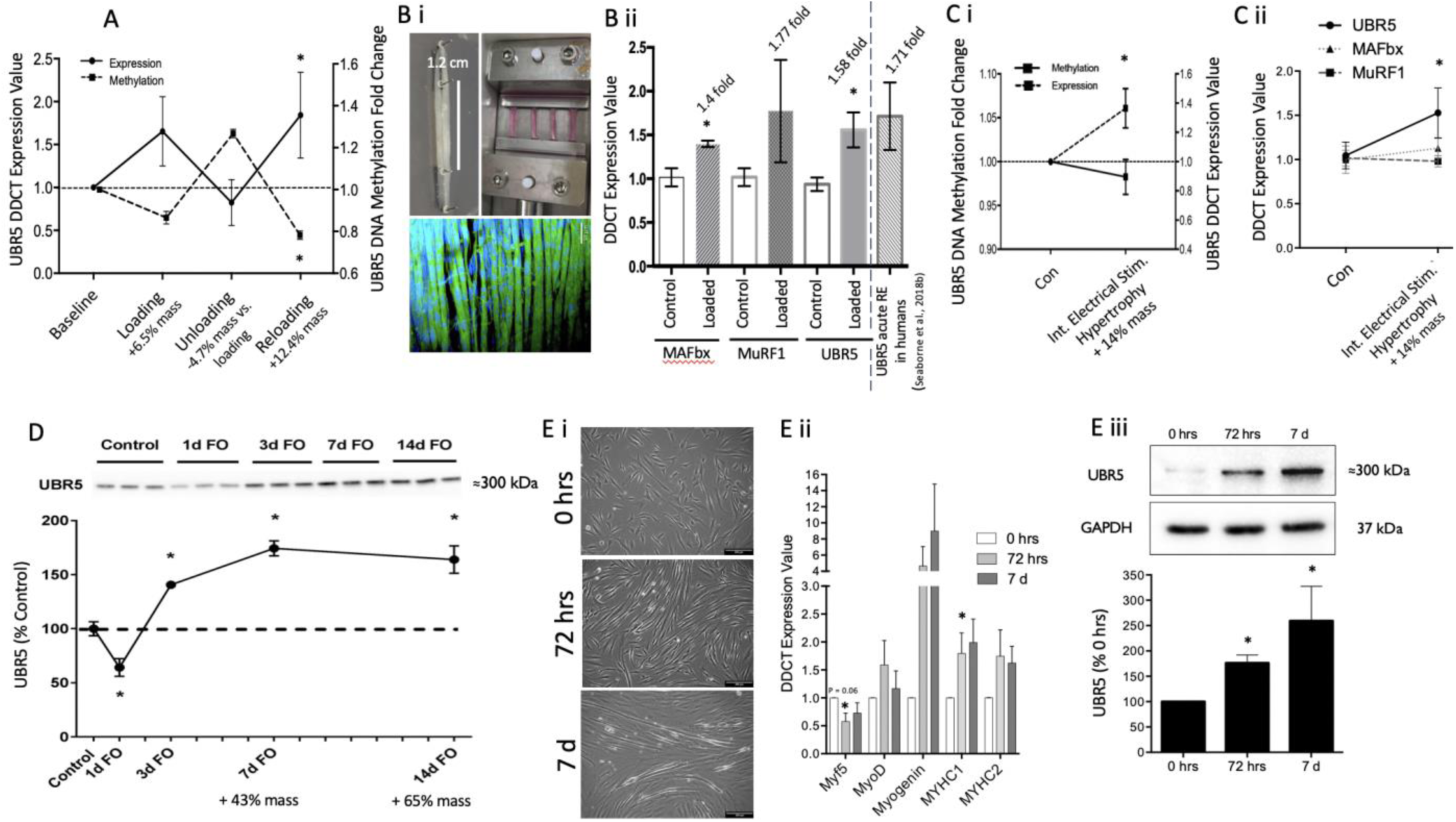
**A:** UBR5 gene expression and DNA methylation redrawn using original data (from (Seaborne *et al.*, 2018b) after an initial period of resistance exercise induced hypertrophy (loading), followed by exercise cessation (unloading) where muscle mass returned to baseline levels, and a later period of resistance exercise induced hypertrophy (reloading). **Bi:** Top image is a representative macroscopic image of c2c12 bioengineered skeletal muscle used as a model to assess UBR5’s response to acute loading. Middle image is a representative microscopic image (middle) of C_2_C_12_ bioengineered skeletal muscle used as a model to assess UBR5’s response to acute loading. Middle image is a representative microscopic image (Nikon, Eclipse Ti-S, 10 × magnification, scale bar = 50 μm) of C_2_C_12_ bioengineered muscle stained with Phalloidin (actin-green) and DAPI (nuclei, blue) demonstrating highly aligned myotubes in the direction of uniaxial tension. Bottom image is the bioengineered muscle constructs loaded within the bioreactor chamber(s) as described in (Turner *et al.*, 2019a). **Bii:** Significant increase in UBR5 gene expression at 30 minutes post-acute loading in bioengineered muscle *in-vitro* (n = 6 loaded / n = 6 unloaded controls), showing similar fold change increase as human tissue after 30 minutes of acute resistance exercise (RE) *in-vivo* in our previous studies (Seaborne *et al.*, 2018b). **Ci & Cii:** UBR5 gene expression significantly increased after a 14% increase in muscle mass as a result of 4 weeks of intermittent high frequency electrical stimulation in rats (n = 4), where UBR5 also demonstrated average reductions in DNA methylation albeit non-significantly (Ci). The increase in UBR5 gene expression in this model of hypertrophy occurred without increases in MuRF1 and MAFbx (Cii). **D:** UBR5 protein abundance significantly reduced in the plantaris muscle in mice after synergist ablation/functional overload (FO) at 1 day (p = 0.02), yet rapidly and significantly increased at 3, 7, and 14 days post FO (all p = 0.001) (n = 3/group). **Ei** Representative images of morphological differentiation of human muscle derived cells in monolayer cultures (n = 3) (scale bar 200 μm), that demonstrated increasing differentiation and myotube formation across the time-course. **Eii:** Myogenic regulatory factors (Myf5, MyoD, Myogenin) and MYHC (I & II) gene expression during human muscle cell differentiation identified an average increase in MyoD, Myogenin and MYHC I & II and reductions in Myf5 as cells differentiated into myotubes at 72 hrs and 7 days. UBR5 gene expression did not increase during differentiation. **Eiii:** UBR5 however demonstrated significant increases in protein levels at 72 hrs and 7 days of differentiation. All graphs are means ± SEM. Significance at the level of p ≤ 0.05 unless otherwise stated in the figure.

We next sought to determine whether the DNA methylation and gene transcript changes observed was related to changes in UBR5 protein abundance using an established model (synergist ablation or functional overload/FO) of load induced hypertrophy of the plantaris muscle in mice. We determined, that after a significant reduction of UBR5 protein 1 d after FO (p = 0.02), UBR5 was increased rapidly and significantly at the protein level at 3, 7 and 14 days over the time-course of hypertrophy (Figure 4D; all p= 0.001) with previously determined corresponding significant increases in muscle weight (43 and 65% at 7 and 14 days respectively, (Baehr *et al.*, 2014)).

Finally, given the differentiation of satellite cells are important for muscle regeneration, and because previous data has described a role for UBR5 in the differentiation of smooth muscle (Hu *et al.*, 2010)), we decided to analyse the temporal gene expression and protein pattern of UBR5 using primary human derived cells over a 7-day time-course of differentiation. Herein we demonstrated that cells increasing in morphological differentiation across the time course of 0, 72 hrs and 7 d (Figure 4Ei), with gene expression increases in MyoD, Myogenin and MYHC2, significant increases in MYHC1 (P = 0.04) and reductions in Myf5 (p = 0.06) (Figure 4E ii), did not demonstrate significant increase in UBR5 at the gene expression level (data Available on request). However, UBR5 protein increased significantly at 72 hrs (p = 0.01) and 7 days (p = 0.02) across the time course of differentiation versus 0 hrs (Figure 4E iii). These data suggest that UBR5 gene expression increased after both acute loading of bioengineered muscle *in-vitro* and chronic intermittent electrical stimulation induced hypertrophy *in-vivo* yet was without changes in mRNA expression during human muscle cell differentiation. Furthermore, UBR5 increases in protein abundance after hypertrophy *in-vivo* and across the time-course of human muscle cell differentiation.

### Genetic Association Studies identified that genetic variants of the UBR5 gene were associated with muscle fiber size in physically active men and strength / power athlete status

Given the increase in gene expression and protein abundance of UBR5 in the above rodent models of hypertrophy, as well as our previous studies in humans demonstrating a strong correlation for UBR5 gene expression with lean leg mass, we tested the hypothesis that genetic variations of the *UBR5* gene might be associated with muscle hypertrophy and strength/power related phenotypes. Firstly, we identified genetic variations in the UBR5 gene that were associated with altered UBR5 gene expression in humans using GTEx (Battle *et al.*, 2017). Using this approach, we identified 13 (rs10110916, rs34695816, rs4302812, rs4621773, rs4734621, rs4734624, rs61504846, rs74195880, rs7814681, rs7844623, rs7845636, rs9693239, rs9694458) single nucleotide polymorphism (SNPs) that were located in close proximity (i.e., in high linkage disequilibrium) near UBR5 gene and were significantly (P≤10^-5^-10^-6^) associated with UBR5 gene expression in human skeletal muscle. Of those, rs4734621 was available for genotyping in the athletic cohorts and controls using micro-array analysis. According to the GTEx, the A allele of the rs4734621 SNP is associated (*P* = 0.000013) with increased expression of the UBR5 gene. It was therefore hypothesized that this allele might be favourable for strength and power-related sports. Indeed, our analysis demonstrated that the rs4734621 AA genotype was associated with high competition results in elite weightlifters (P = 0.016) and the A allele was over-represented in elite strength athletes compared to controls (OR = 1.94, P = 0.04). The rs4734621 A allele was also over-represented in elite power (strength + explosive power) athletes compared to elite endurance athletes (OR = 2.2, P = 0.04). Next, given that the rs4734621 SNP was unavailable to be genotyped in the muscle fibre cohort given the different micro-array platformed used, we identified rs10505025 polymorphism in the UBR5 gene was strongly linked (’D=1) with all 13 SNPs (described above) near the UBR5 gene, including rs473462. We found that the carriers of the rs10505025 AA genotype had significantly (P = 0.024) higher mean (SD) CSA of fast-twitch muscle fibres than carriers of the G allele: AA: 5796 ± 1526 μm^2^; GA: 5260 ± 1350 μm^2^; GG: 4011 ± 983 μm^2^ (Figure 5B). Furthermore, the rs10505025 A allele was significantly over-represented in strength athletes (OR = 2.13, P = 0.05) and sprinters (OR = 2.93, P = 0.03) compared to endurance athletes. These genetic data support our primary findings that the UBR5 gene is associated with skeletal muscle hypertrophy and adaptation to strength/power/resistance training. Further, that variations affecting UBR5 gene expression are associated with cross-sectional area of fast-twitch muscle fibres while favouring strength / power versus endurance status in athletes.

**Figure 5:**
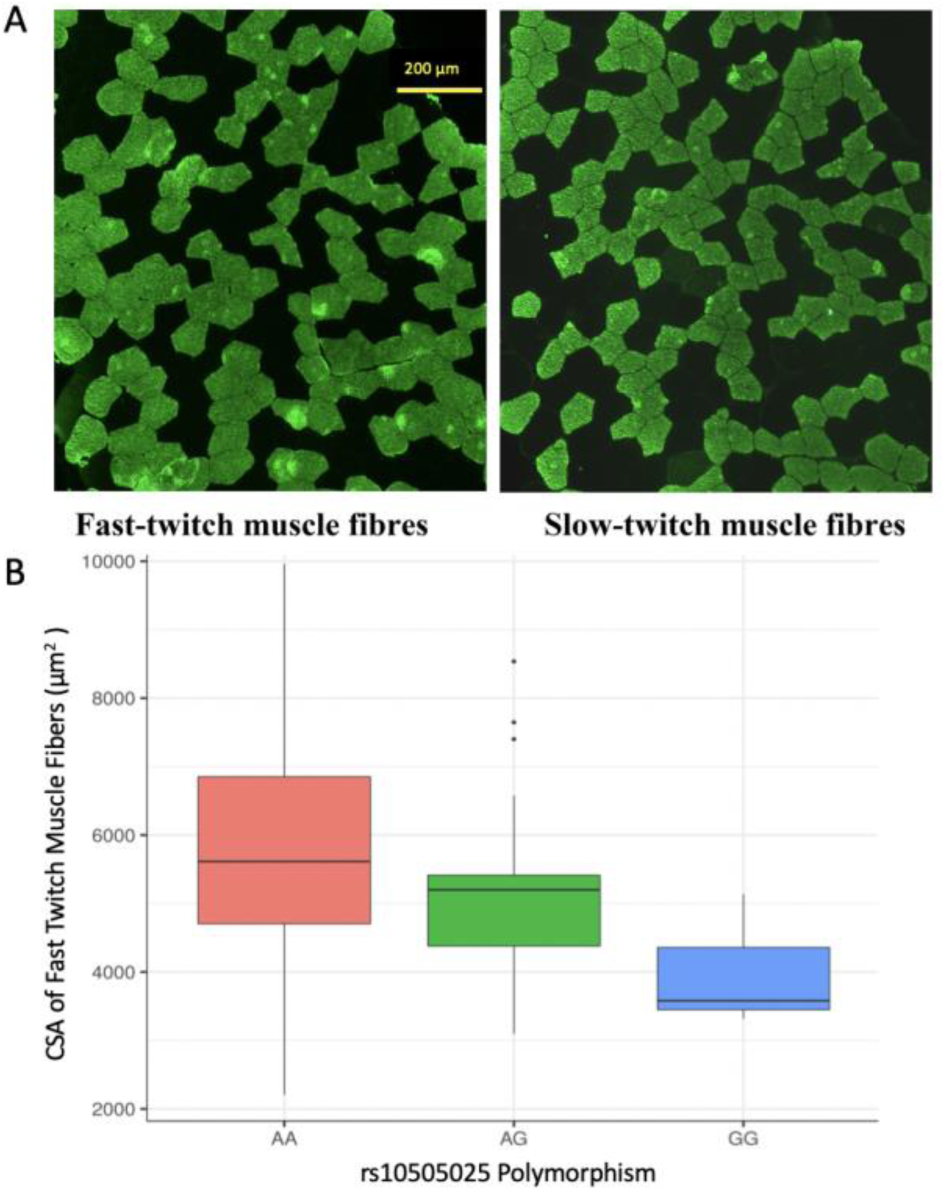
**A**: Example of a slow and fast serial section used for fiber size quantification from human participants. **B**: Box plot demonstrating that there was an increased frequency of large fast fibers within the human population that possessed the AA allele of the UBR5 rs10505025 polymorphism.

## Discussion

### UBR5 in atrophy and recovery from atrophy

UBR5 is a HECT domain E3 ligase previously uncharacterised during and after skeletal muscle atrophy and recovery from atrophy. We demonstrate here that during atrophy evoked by tetrodotoxin (TTX) nerve silencing in rats, the UBR5 promoter was significantly hypomethylated and increased at the gene expression level early (3 days) in the time course of atrophy yet had returned to normal expression levels during later atrophy (14 days) where muscle mass loss was greatest. We have also previously demonstrated that the promoter regions of MuRF1 and MAFbx are hypomethylated during atrophy that corresponds with an increase in gene expression (Fisher *et al.*, 2017), offering support for epigenetic alterations such as DNA methylation being inversely associated with gene expression of other E3 ubiquitin ligases such as UBR5. Furthermore, in the present dataset we provide further support for the alternate roles of UBR5 in comparison with MuRF1 and MAFbx during muscle loss. After 7 days of continuous low frequency electrical stimulation (20Hz) (13% muscle mass loss), MuRF1 and MAFbx were significantly increased while no significant increases were observed in UBR5 expression. Comparable results were obtained after 3 days of atrophy following nerve crush injury, whereby we observed significant increases in UBR5 gene expression that dropped back down at 7 days while MuRF1 and MAFbx remained elevated as muscle continued to waste at 7-14 days. Together, this data supports a potential role for UBR5 in early remodelling after a wasting stimulus such as disuse or nerve silencing rather than in the extensive atrophy of muscle per se.

Because of its potential role in remodelling, we also looked to characterise UBR5 changes during recovery from atrophy as muscle mass returned towards pre-atrophy levels. Indeed, we demonstrated that after 7 days of recovery from TTX induced-atrophy, muscle mass was restored by approximately 52% and UBR5 was significantly hypomethylated and on average gene expression was upregulated while the other E3 ligases, MuRF1 and MAFbx were downregulated. While UBR5 gene expression just failed to reach significance (P = 0.1), this was again interesting given the alternate regulation of UBR5 vs. MuRF1 and MAFbx gene expression. Furthermore, these data support other published observations where MuRF1/MAFbx have been shown not be unchanged in adult rats during recovery from atrophy induced by hindlimb unloading (Baehr *et al.*, 2016). In the TTX model we only observed a partial 52% recovery of TA muscle mass over 7 days, yet saw a significant hypomethylation and average increase in UBR5 gene expression during recovery that might precede any later changes in protein abundance of UBR5. We therefore, measured UBR5 protein levels at later time points post recovery (14 d). Furthermore, due to limited sample from TTX atrophy recovery experiments for analysis of protein (as well as the muscle only being partially recovered 50% at 7 days upon termination of the experiments) we measured protein abundance after atrophy and recovery from hindlimb unloading in rats in new analysis of already published samples, specifically in the medial gastrocnemius muscle that was fully recovered (Baehr *et al.*, 2014). Firstly, we identified that protein levels increased at 3 days following HU but returned to baseline by 14 days where most atrophy occurred, mirroring the temporal regulation observed at the gene expression level in the TTX atrophy model. This suggested that the HU model at the protein level presented a similar response to nerve silencing TTX-atrophy at the gene expression level. Importantly, after 3-and 14-days recovery from HU, we also observed a linear increase in UBR5 protein abundance, with the largest (approx. 50%) increase observed at 14 days where muscle mass was fully recovered. Additionally, there was a more rapid increase in aged muscle during recovery from atrophy at 3 days peaking at an approx. 70% increase at 7 and 14 days during recovery, with these aged animals previously showing impaired recovery vs. young adult rats (Baehr *et al.*, 2014). This suggests that an increase in protein abundance is important during full recovery in healthy young adults, but even a large and robust increase in UBR5 does not enable a full recovery of muscle mass in aged muscle. Finally, after recovery from nerve crush injury in mice, the protein abundance of UBR5 also increased significantly by 50% (similar increase after recovery from HU) and peaked at 28 and 45 days relating to a full recovery of muscle mass after 60 days. Overall, taken together across TTX atrophy, nerve crush and HU models, these data suggest that UBR5 is elevated in response to early atrophy and recovery and supports a role for this gene and protein in remodelling in adult and aged skeletal muscle tissue.

Despite these interesting findings, it is worth mentioning that while UBR5 protein abundance significantly increased at 28 and 45 days following recovery from nerve crush in mice, opposed to the TTX model, UBR5 gene expression did not change. While the reason for no change in mRNA versus increased protein abundance during recovery in the nerve crush model is not clear, it is worth mentioning that we analysed the gastrocnemius muscles because they recovered more over the time course versus the TA (data not shown). Whereas in the TTX model, UBR5 gene expression analysis was in the TA only (given the delivery of the TTX only silences TA and EDL, not gastrocnemius) and the TA is known to have a higher proportion of fast fibres. This discrepancy between gene expression and protein levels following nerve crush in the gastrocnemius versus the other models, leads us to speculate that there are perhaps some important functions for UBR5 at the protein level across different fibre types. However, this requires further empirical evidence to support this hypothesis. Where future experiments should look to perform immunohistochemistry for UBR5 in muscles with different predominant fibre types. Indeed, a similar observation of no increase in gene expression but increases at the protein level occurred during human muscle cell differentiation. Previous studies have suggested UBR5 translocates into the nucleus in a complex that can silence transcription, for example in gene TRAF3 (Cho *et al.*, 2017). However, UBR5 has also been shown to ubiquitinylate Groucho/TLE’s that are repressors of transcription and thus subsequently activate transcription of Armadillo/β-catenin (Flack *et al.*, 2017). Furthermore, there is evidence that UBR5 can regulate translational machinery via interaction with CDK9 (Cojocaru *et al.*, 2011) and PAIP2 (Yoshida *et al.*, 2006). Therefore, this suggests that UBR5 localisation at the protein level, what it is binding to, what it ubiquitinates and its stability/ turnover would be an important avenue of future research in skeletal muscle atrophy and recovery from atrophy. This would be especially prudent in the nerve crush model, given the discrepancy in mRNA and protein data described above. Overall however, the atrophy and recovery data across models combined suggests that UBR5 is elevated early in atrophy and is reduced during later atrophy. Further, UBR5 is elevated during recovery whereas MuRF1/MAFbx are downregulated. More work needs to determine the post-transcriptional regulation of UBR5, its localisation, substrates and binding complexes to further understand its mechanistic role in these processes.

### UBR5’s role in skeletal muscle anabolism and hypertrophy

Given, UBR5’s elevations during early remodelling during atrophy and the subsequent recovery we also aimed to characterise its role in anabolism and hypertrophy. Indeed, the data in this manuscript suggests for the first time that increases in gene expression of the HECT domain E3 ligase, UBR5, occurred following acute loading of mouse bioengineered muscle *in-vitro* and after hypertrophy following chronic intermittent electrical stimulation in rats *in-vivo*. The increase in UBR5 gene expression after acute loading also occurred in the other E3 ligases, MuRF1 and MAFbx, however, after chronic-stimulation induced hypertrophy MuRF1 and MAFbx did not increase whereas UBR5 increased. Together with the data above during atrophy and recovery, this data further supports that, at the gene expression level, UBR5 is alternately regulated versus MuRF1/MAFbx after hypertrophy, but interestingly after acute anabolic loading UBR5, MuRF1 and MAFbx all demonstrate an increase. This is perhaps not surprising given that MuRF1 and MAFbx/Atrogin-1 have also been shown to increase acutely after eccentric contractions in humans (Yang *et al.*, 2006; Louis *et al.*, 2007). Furthermore, the increase in UBR5 gene expression after loading in bioengineered muscle (1.58 fold) supports our previous work in humans, demonstrating a similar fold change in UBR5 gene expression (1.71 fold) after a single bout of resistance exercise in humans (Seaborne *et al.*, 2018b). While gene expression was significantly elevated, the present study identified only 1 out of 6 CpG sites located in UBR5 promoter to be hypomethylated after acute loading in bioengineered muscle, however this was not significant. This supported our previous studies where there was no change in UBR5 DNA methylation after acute resistance exercise in humans (Seaborne *et al.*, 2018b). Despite this, significant hypomethylation of UBR5 has been demonstrated to occur after repeated/chronic training and retraining induced hypertrophy in humans (Seaborne *et al.*, 2018b) and in the present study hypertrophy (14% increase in mass) in rat muscle after chronic intermittent electrical simulation evoked mean reductions in promoter UBR5 methylation (hypomethylation). However, in contrast with the aforementioned human study, the changes did not reach statistical significance. Currently, the data points towards repeated and chronic loading/exercise stimuli being required to cause promoter UBR5 hypomethylation, yet gene expression of UBR5 is elevated in response to both acute and chronic exercise stimuli. Also, given our previous work suggesting that hypomethylation of UBR5 is retained during detraining after a period of training induced atrophy, it is possible that once hypomethylation occurs it is relatively stable, and these changes can be retained over longer periods even when hypertrophic stimuli are removed. This hypothesis requires further investigation to be confirmed.

In addition to significantly increased gene expression after acute anabolic and chronic hypertrophic stimuli in the present study, we also demonstrate that UBR5 protein abundance increased during human muscle cell differentiation and after hypertrophy *in-vivo* following synergist ablation (functional overload/FO) in mice. Indeed, during FO at first there was a significant decrease in UBR5 at 1 d, followed by a rapid and large increase at 3, 7 and 14 days corresponding with extensive hypertrophy of 65% increase in muscle mass by 14 days. It is worth noting that while the initial significant reduction in UBR5 protein at 1d was initially surprising, in the same experiments and samples in earlier published studies at the gene expression level, MuRF1 and MAFbx mRNA were oppositely regulated (elevated) (Baehr *et al.*, 2014), however were then downregulated at 3,7 and 14 days. This again suggesting that MuRF/MAFbx have alternate regulation compared to UBR5, as our other data confirms above across the different models of atrophy, recovery and hypertrophy. Overall, this is the first study to associate increases in protein abundance of UBR5 with differentiation and hypertrophy in skeletal muscle. Future studies should look to overexpress UBR5 in differentiating skeletal muscle cells and also in tissue *in-vivo* to confirm its role in hypertrophy. Furthermore, because UBR5 is an E3 ligase, with apparently alternate regulation to the more extensively studied MuRF1 and MAFbx at the expression level, future studies should look to investigate its protein substrates. For example, studies in cancer cells have suggested that UBR5 can destabilise tumour suppressors such as MDM2/p53 by ubiquitination of upstream ribosomal protein L23 (RPL23), as well as tumour suppressor ECRG4, and its overexpression therefore enables growth and a reduction of apoptosis in colorectal cancer cells (Wang *et al.*, 2017; Watanabe *et al.*, 2018). Therefore, these interactions warrant further investigation in models of skeletal muscle anabolism and hypertrophy. Other studies also suggest that UBR5’s protein activity is important, due to phosphorylation by p90RSK (Cho *et al.*, 2017) and MAPK1 (ERK2) in cells in response to epidermal growth factor (EGF) (Eblen *et al.*, 2003), where ERK is a well-established regulator of muscle cell proliferation (Foulstone *et al.*, 2004) and load-induced hypertrophy (Miyazaki *et al.*, 2011) and p90RSK for protein synthesis in skeletal muscle tissue (Moore *et al.*, 2011). Furthermore, in cancer, overexpression of UBR5 promotes tumour growth via destabilisation of PTEN (an inhibitor of PI3K/Akt) therefore allowing signalling via PI3K/AKT (Zhang *et al.*, 2019), where PTEN and PI3K/Akt are key regulators of both muscle cell differentiation (Foulstone *et al.*, 2004; Sharples *et al.*, 2013) and muscle protein synthesis (reviewed in (Wackerhage *et al.*, 2019)). Finally, UBR5 has been shown to destabilize p21 (Ji *et al.*, 2017), where low p21 is associated with maintenance of muscle during periods of atrophy and high levels of p21 associated with atrophy itself (Bongers *et al.*, 2015). Therefore, UBR5 signalling (ERK, PTEN/PI3K/Akt/p90RSK), its targets (RPL23/p53, ECRG4, p21) and cellular location/transcription factor binding (Groucho/TLE’s, TRAF3) will be key future avenues of research to uncover its mechanistic role in muscle cell differentiation and muscle hypertrophy.

Finally, we investigated genetic variations of the UBR5 gene that were associated with UBR5 gene expression in humans using the online tool GTEx (Battle *et al.*, 2017) to further corroborate the animal *in vivo* data, human *in vitro* data and previous *in vivo* human data (Seaborne *et al.*, 2018b). Subsequently, the genetic variants that were identified were found significant in various human and athlete cohorts including physically active men, strength/power or endurance trained athletes. Indeed, we identified 2 SNPs, the first rs4734621 AA genotype (associated with increased UBR5 gene expression) was related to high competition results in elite weightlifters and the rs4734621 A allele was over-represented in both elite strength athletes versus controls and elite power athletes versus endurance athletes. Further, the A allele of the rs10505025 polymorphism (which is strongly linked with rs4734621) was associated with greater CSA of fast-twitch muscle fibres in physically active men and was significantly over-represented in strength athletes and sprinters compared to endurance athletes. These genetic data support the animal data suggesting that where an increase in gene expression of UBR5 is expected that this is associated with hypertrophy. Furthermore, that adaptation to strength/resistance exercise and muscle size of fast fibres is associated with the aforementioned UBR5 polymorphisms in humans. The only study to date to investigate other E3 ligases polymorphisms and human muscle are studies in MuRF1 (rs2275950) that describe the AA homozygotes were stronger and demonstrate less muscle soreness in response to eccentric muscle damage compared with GG homozygotes (Baumert *et al.*, 2018). At present no other studies have identified UBR5 polymorphisms to be associated with skeletal muscle size and performance in humans.

## Conclusion

Using multiple models of skeletal muscle atrophy/injury, recovery from atrophy, anabolism and hypertrophy *in-vivo* and *in-vitro*, we demonstrate for the first time that the E3 ubiquitin ligase UBR5, in alternate fashion to MuRF1/MAFbx at the gene expression level, is elevated during recovery, anabolism and hypertrophy in animals *in-vivo* as well as human muscle cell regeneration *in-vitro*. Finally, in humans the A alleles of the rs10505025 and rs4734621 SNPs that affect the expression of the *UBR5* gene are strongly associated with larger fast-twitch muscle fibres and strength/power performance.

## Acknowledgments

These data were supported by GlaxoSmithKline, Society for Endocrinology, North Staffordshire Medical Institute (NMSI) awarded to APS (PI) and UK Engineering and Physical Sciences Research Council (EPSRC) / UK Medical Research Council (MRC) centre for doctoral training studentship awarded to PG in the group of APS (PI). Funds from the Doctoral Alliance / Liverpool John Moores and Keele University PhD scholarships underpinned the work by RAS and DCT via APS. Rodent TTX and electrical stimulation work was supported by an integrative mammalian biology studentship from the UK Medical Research Council (MRC) awarded to the group of JCJ (PI). The human genetics study was supported in part by grant from the Russian Science Foundation, Grant No. 17-15-01436. We would also like to thank: Elena S. Kostryukova, Nickolay A. Kulemin, Sofya A. Khabibova, Alexander V. Pavlenko, Ekaterina V. Lyubaeva, Evgeny A. Lysenko, Tatiana F. Vepkhvadze and Egor M. Lednev for all their work/input on the human genetics study.

## Author contributions

APS conceived research. APS, SCB, JCJ, IIA, EVG, DJO, DCH designed the research. APS, SCB, JCJ, DCT, DCH, RAS, DJO, IIA, LMB, PG, EAS, OVB, AKL, DVP, EVG authors performed the research. All authors were involved in data analysis. APS drafted the manuscript. All authors reviewed the manuscript drafts and inputted corrections, amendments and their expertise.

